# SCS: cell segmentation for high-resolution spatial transcriptomics

**DOI:** 10.1101/2023.01.11.523658

**Authors:** Hao Chen, Dongshunyi Li, Ziv Bar-Joseph

## Abstract

Spatial transcriptomics promises to greatly improve our understanding of tissue organization and cell-cell interactions. While most current platforms for spatial transcriptomics only offer multi-cellular resolution, with 10-15 cells per spot, recent technologies provide a much denser spot placement leading to sub-cellular resolution. A key challenge for these newer methods is cell segmentation and the assignment of spots to cells. Traditional image-based segmentation methods are limited and do not make full use of the information profiled by spatial transcriptomics. Here we present SCS, which combines imaging data with sequencing data to improve cell segmentation accuracy. SCS assigns spots to cells by adaptively learning the position of each spot relative to the center of its cell using a transformer neural network. SCS was tested on two new sub-cellular spatial transcriptomics technologies and outperformed traditional image-based segmentation methods. SCS achieved better accuracy, identified more cells, and provided more realistic cell size estimation. Sub-cellular analysis of RNAs using SCS spots assignments provides information on RNA localization and further supports the segmentation results.

## Introduction

Spatial transcriptomics provides information on both the expression and the locations of cells in a sample, enabling the analyses of cell-cell signaling and cell type organization [1, 2]. Spatial transcriptomics uses a set of barcoded spots placed at regular intervals on the sample to profile the expression of genes [3]. To date, most platforms (including Visium [3]) use spots that were placed up to 100 *µ*m apart. In such placement, each spot captures RNAs from 10-20 cells depending on the tissue, making it hard for this technology to achieve single cell-level analysis. Very recently, new spatial technologies enable a much denser spot placement. For example, both Stereo-seq [4] and Seq-scope [5] achieve a spot-to-spot distance of 0.5 *µ*m on average, resulting in more than 1000 spots per cell on average.

While promising, a challenging step in the analysis of such new data is cell segmentation and the integration of spots to assign an expression profile to each cell. Recently, new segmentation methods were developed for spatial proteomics [6] or *in situ* fluorescent hybridization (FISH) based spatial transcriptomics data [7, 8]. For example, Baysor [7] spatially clusters observed molecules to cells for FISH based data by modeling each cell as an ellipsoid using Gaussian distribution of transcript composition. However, the detected molecules in such data are not restricted to predefined spot locations, which makes it much easier to model cell shapes. In addition, the large number of genes profiled and the sparseness of the expression captured by each spot make these methods not appropriate for sequencing based technologies. Most standard cell segmentation methods developed to date rely on nucleus or membrane straining to identify cell boundaries [9, 10, 11, 12, 13]. While successful, these methods do not fully utilize the information provided by spatial transcriptomics data, leading to less accurate results. In addition, most of them [10, 11, 12, 13] require manual annotations for model training, which are usually not easy to obtain in sufficient quantities when profiling new tissues.

To address these issues, we developed SCS (Sub-cellular spatial transcriptomics Cell Segmentation), which combines sequencing and staining data to improve cell segmentation in high-resolution spatial transcriptomics. SCS performs segmentation in three key steps (Fig. 1). It first identifies cell nuclei from staining images using the Watershed algorithm [9]. Second, a transformer model infers for each spot whether it is part of a cell or part of the extracellular matrix (background), and its relative position w.r.t. the center of its cell, by adaptively aggregating high-dimensional but sparse gene expression information from neighboring spots via an attention mechanism. To train the model, we used as positive examples spots within the identified nuclei and as negative samples spots sampled from highly confident background regions. Finally, spots that are determined to be part of the cell are grouped by tracking the gradient flow from spots to nucleus centers. See Methods for complete details.

**Figure 1:**
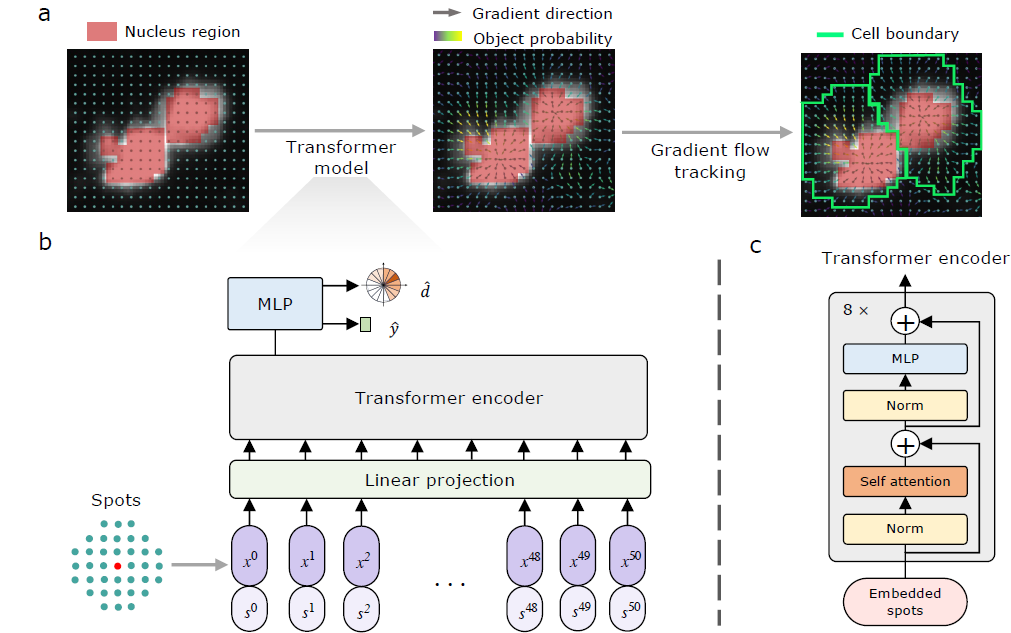
Workflow of SCS. **a**, Barcoded spots (cyan dots) that reside inside a cell nuclei (red masks) are first identified by segmenting the stained image. A transformer model is next trained on these spots and some background spots to predict the gradient direction (arrow) from each spot to the center of the cell to which it belongs and the probability that it is part of a cell (yellow arrow) or part of the extracellular matrix (purple arrow). The transformer model is then applied to all other spots. A gradient flow tracking algorithm is used to segment cells by grouping spots based on their gradient prediction. **b**, The transformer model predicts for each input spot the probabilities from this spot to its cell center for 16 predefined directions (*d̂*) and the probability that the spot is part of a cell (*ŷ*). For each spot (red dot), the transformer model aggregates information from its 50 nearest neighboring spots (cyan dots) by adaptively learning a weighting based on the spot expression (*x*) and relative positions (*s*). **c**, The structure of one transformer encoder layer, see “Methods” for details.

We applied SCS to two different high-resolution spatial transcriptomics platforms: a mouse brain dataset profiled using Stereo-seq [4] accompanied with nucleus staining, and a mouse liver dataset from Seq-scope [5] accompanied with H&E staining. To evaluate performance, we compared SCS with Watershed cell segmentation, which is used by these platforms, and popular deep learning based segmentation methods, including Cellpose [10], DeepCell [11], and StarDist [12]. As we show, our method obtained more accurate segmentation, more cells, and more realistic cell sizes when compared to these methods. In addition, we further used SCS to analyze the sub-cellular localization of different RNAs and show that our results agree with prior knowledge further validating the accuracy of our segmentation.

## Results

### Application of SCS to high-resolution spatial transcriptomics data

We tested SCS on public subcellular-resolution *in situ* datasets generated using two different platforms: a mouse brain dataset profiling using Stereo-seq [4] and a mouse liver dataset that utilized Seq-scope [5].

The Stereo-seq dataset captures a whole adult mouse brain slice (an area of around 5.3 mm×7.0 mm) in a single section. The barcoded spots are arranged in a grid with a distance of 0.5 *µ*m between spots (which, given an average cell size of 20 *µ*m in diameter means that there are roughly 1200 spots per cell, Extended Data Fig. 1). In total, this dataset profiled 26,177 genes in more than 42,000,000 spots with an average of 3.3 unique molecular identifier (UMI) counts per spot (Extended Data Fig. 2). The brain slice was imaged with nucleic acid staining, allowing for segmentation of the nucleus using image based methods.

**Figure 2:**
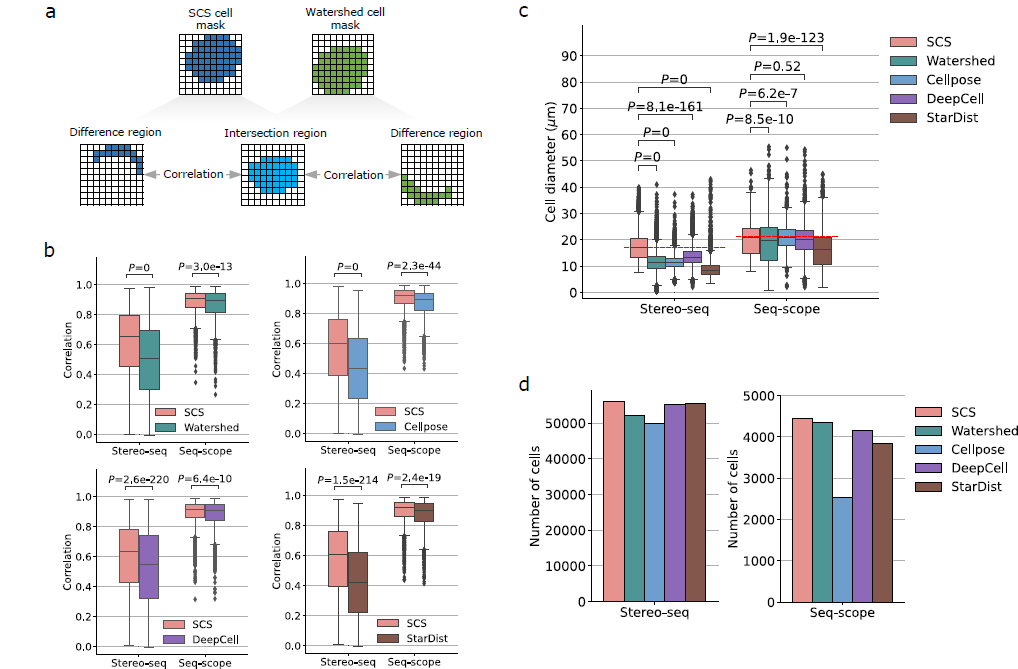
Performance evaluation of SCS and comparisons with other methods. **a**, The benchmark used for evaluating the performance and for comparison of different segmentation methods. The intersection region and respective difference (unique) regions between two segmentation are calculated for each cell. The segmentation is said to have higher accuracy if its unique cell region is better correlated with the intersection region. **b**, Comparison between SCS and the four other image segmentation methods using the correlation benchmark. Each cell used for evaluation contributes one point to the correlation presented in the boxplot. SCS achieved significantly higher segmentation accuracy than other methods on both datasets (Wilcoxon signed-rank tests, one-sided; for SCS vs. Watershed, *N* =17,811 cells on Stereo-seq, *N* =2,757 cells on Seq-scope; for SCS vs. Cellpose, *N* =13,310 cells on Stereo-seq, *N* =2,446 cells on Seq-scope; for SCS vs. DeepCell, *N* =26,916 cells on Stereo-seq, *N* =3,513 cells on Seq-scope, for SCS vs. StarDist, *N* =6,370 cells on Stereo-seq, *N* =2,635 cells on Seq-scope. Supplementary Note 3 for cell filtering criteria). **c**, Comparison of the sizes of cells segmented by SCS and other methods. SCS obtained segmented cells with larger cell diameters than all the other methods with significant differences on Stereo-seq (Kruskal-Wallis tests; for Stereo-seq, *N* =56,187 for SCS, *N* =52,004 for Watershed, *N* =50,020 for Cellpose, *N* =55,260 for DeepCell, *N* =55,364 for StarDist; for Seq-scope, *N* =4,456 for SCS, *N* =4,354 for Watershed, *N* =2,527 for Cellpose, *N* =4,157 for DeepCell, *N* =3,832 for StarDist). The red dashed lines show the expected cell diameters from the literature. **d**, The number of cells identified by the segmentation methods for the two datasets. **b,c**, Boxplots show medians (horizon-tal line in each box), interquartile ranges (boxes), 1.5 interquartile (whiskers), and remaining point individuals.

The Seq-scope dataset contains four tissue sections from mouse liver. Each section captures around a 0.93 mm×0.80 mm region of the tissue. The center-to-center distance of barcoded spots is similar to the distance for the Stereo-seq data, 0.5 *µ*m on average. In total, 24,171 genes were profiled in four sections and each section contains over 570,000 spots (Extended Data Fig. 3), with 5.7 UMI counts per spot on average (Extended Data Fig. 4). Instead of using nuclei staining, the Seq-scope protocol images tissues with the hematoxylin and eosin (H&E) staining. Therefore, the entire cell bodies can be segmented using the imaging data.

**Figure 3:**
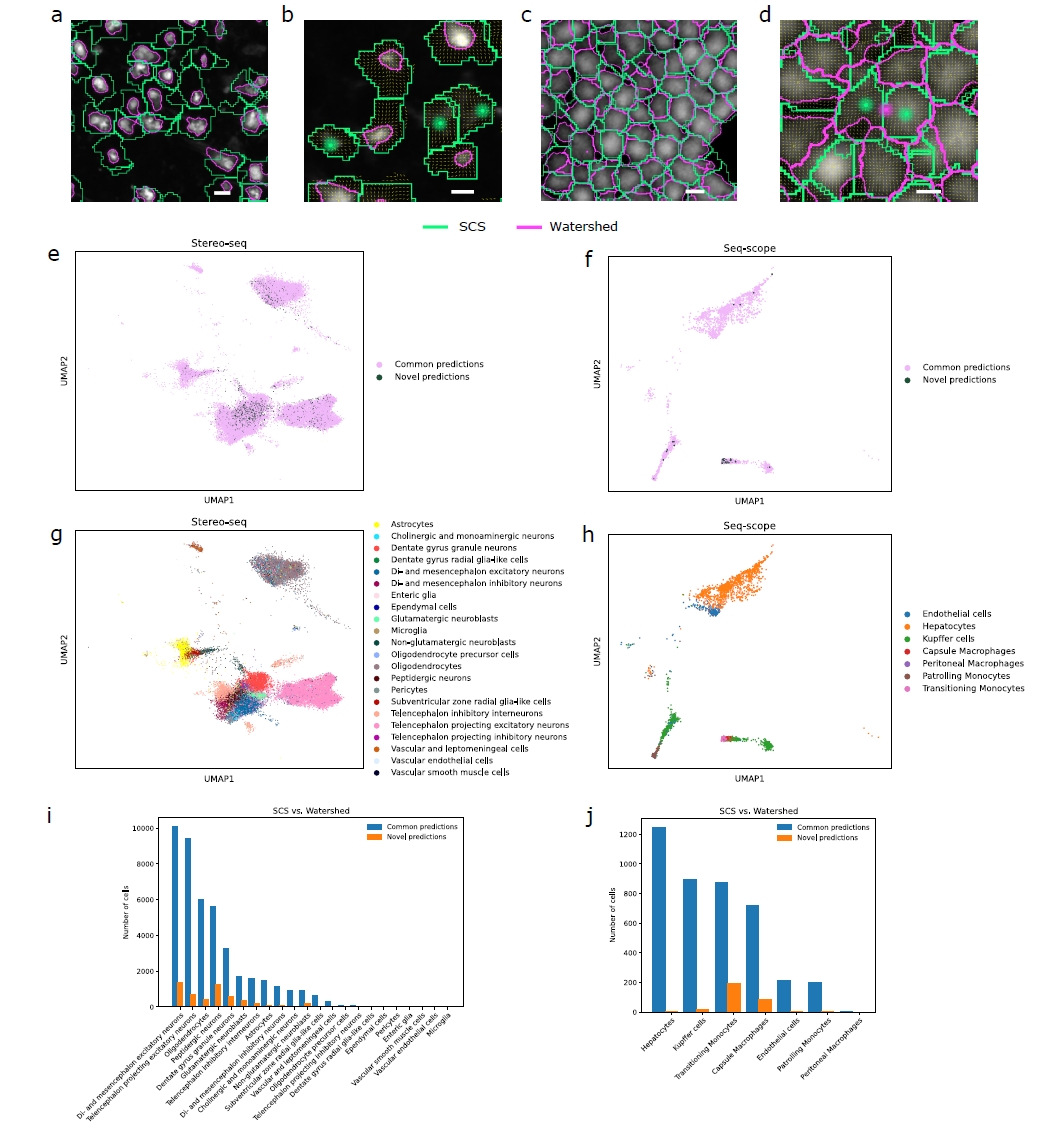
Cell segmentation examples and the distributions of cells in low dimensional space. **a**, SCS captured cytoplasm regions of cells in the nucleus staining and thus segmented cells with larger sizes. **b**, Example of segmentation results on the Stereo-seq dataset where Watershed missed three cells due to their low staining signal intensity while SCS identified them (green dots). **c**, Two segmentations show similar cell sizes but with disagreement on cell boundaries on the Seq-scope dataset. **d**, Segmentation example on the Seq-scope dataset where Watershed merged two cells as one cell (pink dot) due to their unclear boundary in the image while SCS successfully segmented them (green dots). **e**, UMAP projection of SCS segmented cells on the Stereo-seq dataset based on their expression profiles. Novel predictions (darker nodes) are mixed with cells identified using image segmentation. **f**, UMAP projection of SCS segmented cells on the Seq-scope dataset. **g**, Cell type annotation for the Stereo-seq dataset. **h**, Cell type annotation for the Seq-scope dataset. **i**, The number of novel cell predictions by SCS compared to Watershed vs. the number of cells that are commonly identified by SCS and Watershed in different cell types for the Stereo-seq dataset. Cell types with more novel predictions are usually those with smaller nucleus sizes as shown in Supplementary Figure 4a. **j**, The same comparisons for the Seq-scope dataset. **a-d**, Experiments that generated the examples were independently repeated three times with similar results. Scale bars: 10 *µ*m.

**Figure 4:**
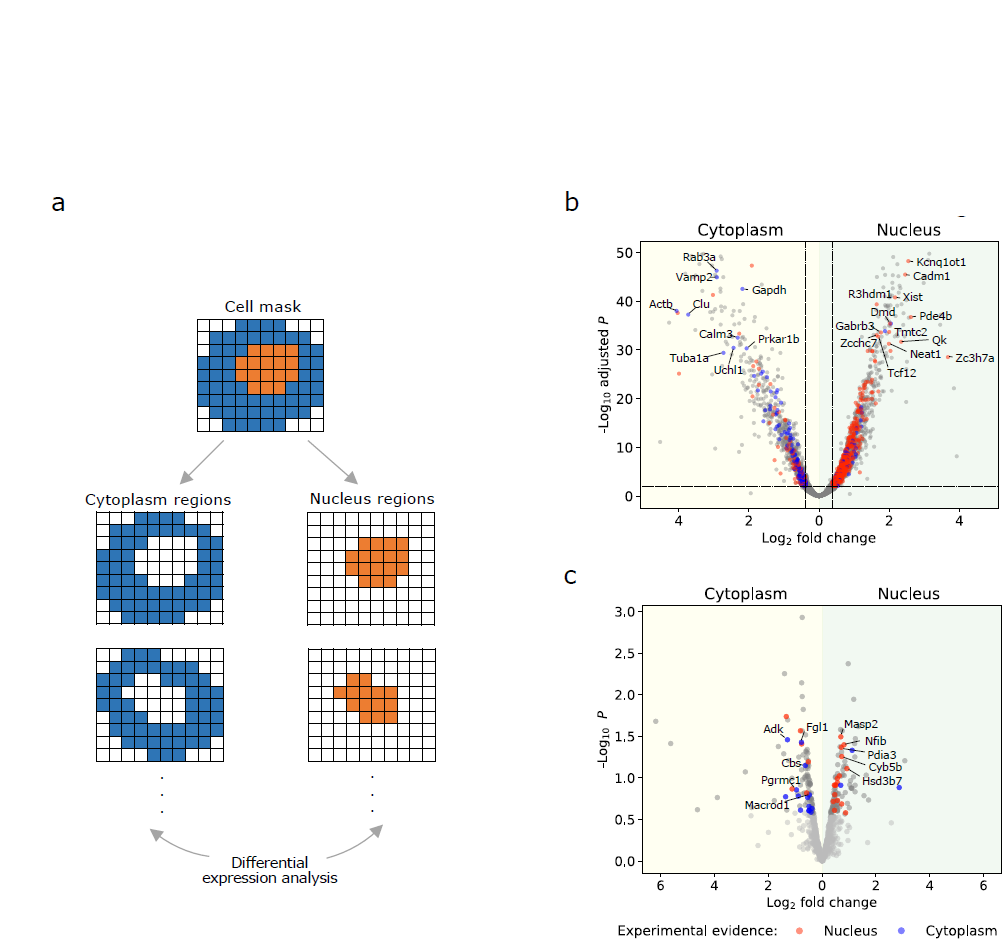
Sub-cellular analysis using SCS. **a**, Identification of genes whose RNAs are differ-entially localized. **b**, Volcano plot that shows quantitative changes in expression levels for genes between the nucleus and cytoplasm of SCS segmented cells for the Stereo-seq dataset. Genes with *P*-values *<* 0.01 and fold changes greater than 1.3 were identified from each group (t-tests, one-sided, the Benjamini-hochberg method was used to adjust *P*-values; *N* =31,763 nucleus regions, *N* =37,940 cytoplasm regions, subcellular regions with at least 100 genes were used for this analysis). Genes whose RNAs have been experimentally shown to reside in the nucleus or cytoplasm are colored accordingly. **c**, Volcano plot for the Seq-scope dataset. The top 100 genes with the smallest *P*-values were identified from each group (t-tests, one-sided, raw *P*-values are shown to avoid most of the *P*-values being corrected to the same value; *N* =2,779 nucleus regions, *N* =3,006 cytoplasm regions, regions with at least 100 genes were used for this analysis).

### SCS provides accurate cell segmentation

We first applied SCS to the Stereo-seq data. To evaluate its performance, we compared SCS with Watershed cell segmentation (Supplementary Note 1), and other popular segmentation methods that are based on deep learning including Cellpose [10], DeepCell [11], and StarDist [12]. Appropriate pretrained models of the deep learning methods are used for evaluation (Supplementary Note 2). Since ground truth for cell segmentation does not exist, we used a popular method for evaluating cell segmentation methods [6, 7]. In this evaluation, we compare the expression of regions where two methods (SCS and another method) agree to regions where they disagree (Fig. 2a). Specifically, for each nucleus, we identified a cell mask using the segmentation of each method. The intersection and difference regions between the two cell masks for this nucleus were then computed. We next estimated the correlation of expression profiles between the intersection region and each of the difference regions. Since the intersection region is often dominated by the nucleus, which is easily detected by all methods (much easier to stain), we treat it as ground truth and compare the non-intersecting regions to the intersection. We expect that the more correlated the difference region is with the intersection region the more accurate the segmentation of the method.

On the Stereo-seq dataset, SCS segmentation achieved an average correlation 24% higher than that of Watershed (0.61 vs. 0.49) (Fig. 2b), and at least 13% higher than those of all other deep learning segmentation methods (0.60 vs. 0.53 of DeepCell). While image based methods on nucleus staining images tend to underestimate cell sizes, SCS is able to accurately capture cytoplasm regions of cells (Fig. 3a). As a result, the segmented cells of SCS show more realistic cell diameter [14] compared to other segmentation methods (Fig. 2c, Supplementary Note 4). We further observed that some cells were completely missed by image based method due to their low staining signal intensity. However, SCS can identify such cells based solely on transcriptomics data (“Novel predictions”, Fig. 3b, Supplementary Note 5). On the other hand, the cells only identified by image based methods but missed by SCS are much fewer (Supplementary Fig. 1), leading to at least 1.5% more cells of SCS than all the other methods (56,187 vs. 55,364 of StarDist which is the 2nd highest, Fig. 2d). The UMAP [15] projection of SCS segmented cells based on their expression profiles shows that these novel predictions match cells using image segmentation (Fig. 3e), indicating that they are likely real cells. The staining signals for the regions covered by the novel cell predictions also support the accuracy of these novel predictions (Supplementary Fig. 2).

For the Seq-scope data, the differences were less dramatic due to the use of H&E images. Still, SCS had a higher correlation of 0.88 vs. 0.86 for Watershed and also higher correlations than all the other deep learning based methods (Fig. 2b). All methods achieved higher correlations when using cell stained images when compared to nucleus staining, as expected (although, as we show, by using the expression values SCS can still improve for this type of staining as well). As the cell stained images were used, different methods show similar cell sizes. SCS segmentation leads to cell diameters (19.9 *±* 6.4 *µ*m) that are slightly larger when compared to Watershed (18.3 *±* 9.4 *µ*m) and StarDist (16.2 *±* 7.6 *µ*m) but slightly smaller when compared to Cellpose (21.1 *±* 4.8 *µ*m) and DeepCell (20.1 *±* 6.0 *µ*m) (Fig. 2c and 3c). Again, SCS determined cell size is consistent with previous findings [16]. In addition, we observed that when the boundaries of two cells are unclear in the staining images, image based methods tend to merge them, while SCS can segment them with the help of transcriptomics data (Fig. 3d) leading to at least 2.3% more cells when using SCS segmentation (4,456 vs. 4,354 of Watershed) (Fig. 2d). The UMAP projection shows that the novel cell predictions are mixed with cells that overlap with image based predictions, however mainly correspond to a subset of cell clusters (Fig. 3f). Further investigation by cell type annotation (Supplementary Note 6) indicates that these novel predictions are more likely to be non-parenchymal cells (such as kupffer cells and monocyte-derived cells) than hepatocytes (Fig. 3h, j). Such cells are known to be smaller and so harder to identify using H&E staining [5]. We observe a similar trend when comparing SCS segmentation to other methods as well (Supplementary Figs. 3a-c).

Cell type analysis for the Stereo-seq dataset confirmed the fact that SCS can help recover cells from cell types with smaller cell sizes (Figs. 3g, 3i, Supplementary Fig. 4a). These results hold for all methods we compared to (Supplementary Figs. 3d-f, 4b-d). Spatial analysis of the location of cell types with novel predictions by SCS shows that they are located in areas that are consistent with the relevant histological features of the tissues (Supplementary Figs. 5-6).

Although SCS is designed for high-resolution sequencing based platforms, *in situ* fluorescent hybridization (FISH) based spatial transcriptomics techniques also provide subcellular resolution and are highly multiplexed. We therefore tested SCS on two datasets generated using state-of-the-art FISH based platforms, seqFISH+ [17] and MERFISH [18]. To process the FISH-based data, we transformed detected RNA points to grid-shaped spots (Supplementary Notes 7, 8). Here, in addition to the image-based methods, we also compared SCS with two cell segmentation methods specifically designed for FISH data, Baysor [7] and JSTA [8] (Supplementary Notes 9, 10). Results show that SCS outperforms all image-based segmentation methods on the FISH based data, and outperforms Baysor and JSTA on most of the four datasets we looked at (Supplementary Figs. 7-9). In addition, for the seqFISH+ dataset, manually annotated cell segmentation ground-truth is available [17]. We thus validated the performance of different methods using this data by quantifying the agreement of their segmentation with the ground truth (Intersection over Union (IoU)). SCS achieved a much higher IoU (0.75) than all the other methods (0.47 of Baysor which is the second highest) (Supplementary Fig. 7).

Finally, we also tested DeepCell with the Mesmer model [13] (Supplementary Note 2) and observed that using Mesmer does not improve the DeepCell results (Supplementary Figs. 7-9).

### SCS enables sub-cellular analysis of spatial transcriptomics

While traditional spatial transcriptomics is a powerful method for studying cell type expression and interactions [19, 20], the use of high-resolution methods opens the door to characterizing molecular heterogeneity within individual cells. This can be important to study RNA dynamics and to fully understand cellular variability in tissues [21]. We therefore used SCS to investigate how RNAs are distributed within cells. Specifically, we divided each cell into two regions, the nucleus region (identified using the staining image data) and the cytoplasm region (the rest of the cell mask identified by SCS, Fig. 4a). Genes whose RNAs localize differentially between two groups of regions were identified using t-test (Fig. 4b-c). Interestingly, in both datasets, RNAs that have been experimentally shown to reside in the nucleus or cytoplasm [22] are significantly enriched in our identified RNAs in the corresponding regions (*P* = 4.0 × 10*^-^*^50^ for Stereo-seq, *P* = 6.4 × 10*^-^*^4^ for Seq-scope, Fisher’s exact test, one-sided, Extended Data Fig. 5). For example, the long non-coding RNA (lncRNA) of gene *Kcnq*1*ot*1 is a nuclear transcript that interacts with chromatin and regulates transcription of multiple genes [23]. The lncRNA of gene *Neat*1 is a well-known nuclear transcript that forms the core component of organelles in nucleus [24]. Both RNAs are identified as being differentially localized to nucleus in SCS segmentation (Fig. 4b). In contrast, gene *Rab*3*a* and gene *V amp*2 both encode proteins that are involved in neurotransmitter release and associated with cytoplasmic vesicles [25, 26]. They are both found with high expression levels in the cytoplasm regions in SCS segmentation (Fig. 4b).

These experiments were repeated for the other segmentation methods we compared to. We found that for all other methods, we obtained fewer RNAs that are experimentally verified when compared to those identified by SCS (Supplementary Fig. 10). These results indicate that SCS segmentation can better help sub-cellular analysis in high-resolution spatial transcriptomics data and partially explains why our model can obtain better cell segmentation with transcriptomics data.

## Discussion

A key step in the analysis of spatial transcriptomics data is cell segmentation. This is especially true when using the very recent sub-cellular profiling platforms. For such data, accurate cell segmentation is essential as errors in identifying cell boundaries can directly impact gene expression level quantification in cells, and further influence downstream analysis. Existing cell segmentation methods for this data only rely on the stained image, which do not fully utilize the information provided by the experiment leading to less accurate results.

In this study, we developed a cell segmentation method that combines both the staining information and the expression data to refine cell segmentation. Unlike prior methods, our method focuses on the spots but not the staining. For each spot, the method attempts to determine whether it is within a cell and if so, which cell. Once such assignments are determined, cells are naturally segmented by grouping all the spots that belong to the same cell. To enable spot assignment, SCS first aggregates information from neighboring spots and then maps spots to low-dimensional latent representations to determine their relative positions to the centers of cells. To train the supervised model, we first identify the nucleus regions which serve as ground truth and then use as positive examples spots in these regions and as negative samples spots sampled from highly confident background regions to train the transformer model, which is then applied to the whole section.

Applications of SCS to two datasets generated using very recent state-of-the-art *in situ* capturing platforms demonstrates the advantage of our method. SCS segmentation achieves higher segmentation accuracy, detects more cells, and yields more realistic cell sizes when compared to several widely used image based segmentation methods. Analysis of the spatial distribution of RNAs identified many RNAs enriched in different sub-cellular regions and these agree with experimentally confirmed results. These findings further validate SCS segmentation and suggest the ability of SCS in facilitating sub-cellular analysis on high-resolution spatial transcriptomics.

Although SCS works well for the platforms we tested, there are a number of ways to further improve it. Cell shape information can be further utilized to better obtain cell masks given the high-density spot placement. In addition, feature selection can be improved. In this study, we used the 2,000 variable genes as the input of the transformer model though better results may be obtained by a more sophisticated feature selection method. Moreover, if RNA spatial distribution patterns are found to vary by cell types in the datasets [27], cell-type specific attention layers can be introduced (based on the expression profile) to allow the model to learn different patterns and better segment cells. Finally, while the integration of high-dimensional transcriptomics and imaging data is computationally intensive, such runtime is still reasonable for the analysis of this complex data (Supplementary Table 1). In addition, runtime and memory can be further optimized for specific platforms.

Since ground truth is generally unavailable for most spatial transcriptomics data, finer-grained cell type analysis might provide more evidence for cell segmentation quality. For example, a method that can recover more reasonable cell type distributions or detect rare cell types missed by other methods might be more powerful. We have performed some preliminary analysis of this in the paper (Supplementary Fig. 11) but more detailed analysis may be required to fully validate this point.

SCS was implemented in Python and is available for download at: https://github.com/chenhcs/SCS. While sub-cellular spatial transcriptomics is still very new, we believe that its advantages and ability to provide spatially resolved single cell information would make it very popular going forward. We hope that SCS would be a useful pre-processing method to enable all downstream analysis of such data.

## Acknowledgements

We thank Dr. Longqi Liu, Dr. Jun Hee Lee and Dr. Rongxin Fang for sharing the Stereo-seq, Seq-scope and MERFISH data, respectively, and for advising us on how to process these datasets. Work partially supported by National Institutes of Health (NIH) grants OT2OD026682, 1U54AG075931 and 1U24CA268108 to Z.B.-J.

## Author Contributions

H.C., D.L., and Z.B.-J. conceptualized and designed the study. H.C., D.L., and Z.B.-J. designed the algorithm and methodology. H.C. developed the software of SCS with supervision from Z.B.-J. H.C. and D.L. performed evaluations and result analyses. H.C., D.L., and Z.B.-J. wrote the manuscript. All authors read and approved the final manuscript.

## Methods

### Data preprocessing

Gene counts in each barcoded spot were collected from the original paper of Stereo-seq [1] and Seq-scope [2], and used to generate a gene expression profile vector for each spot. Each element in the profile represents the number of transcripts observed in the spot for that gene.

To identify nuclei, the paired staining image and sequencing section are first aligned to match image pixels and spot coordinates. For this, a count heat map was created for the sequencing section, where each element in the heat map contains the total number of detected transcripts in a spot. The staining image was then aligned to the heat map using transformations implemented in Spateo (https://spateo-release.readthedocs.io) [1]. This step was omitted for the Seq-scope dataset as the images have been prealigned. Watershed algorithm implemented in Spateo was next used to segment nuclei from the aligned staining image, and the mask of each individual nucleus was obtained (Supplementary Note 1).

### Transformer model for spot-level predictions

An overview of the model is depicted in Figure 1b. The model contains two components: an encoder and a classifier. The encoder maps each spot to a hidden representation *z*. Given the very sparse set of genes that are usually detected for each spot, we combine neighborhood gene expression information when generating the hidden representation for a spot as follows. For each spot, we use its expression profile and combine it with the profiles of the 50 nearest neighbor spots, represented as *x* = (*x*^0^*, x*^1^*, …, x*^50^). Each vector *x^i^* has *N* dimensions which represent the number of detected transcripts in that spot for each of the *N* genes used in the study. The nearest neighbors are defined by using the euclidean distance between spot coordinates. To enable the model to use the relative locations of different neighbors, we include in the input to the encoder the distance from the center spot to each of the neighbor spots denoted as *s* = (*s*^0^*, s*^1^*, …, s*^50^), where *s^i^* is a two dimensional vector containing distances on two axes. Given the resulting hidden vector *z* for a spot, a classifier then predicts the direction from the spot to the center of its cell denoted by *d̂*, and the probability it belongs to part of a cell, *ŷ*.

### Encoder

The expression profiles, *x*, and the distance vectors, *s*, for the center spot and neighbor spots are first projected to a set of *D* = 64 dimensional vectors *r* through two separate fully connected layers followed by summation:

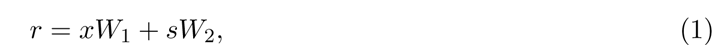

where *W*_1_ ∈ ℝ *^N×D^* and *W*_2_ ∈ ℝ ^2×*D*^ are weights matrices of the two dense layers.

We used a similar network architecture as the one proposed in the original Transformer paper [3]. In this architecture, the encoder is composed of a stack of *L* = 8 identical layers. Each layer has two blocks. The first is a self-attention block (SA):

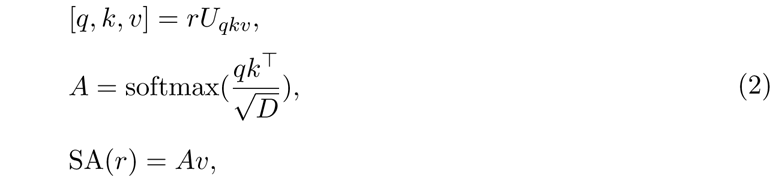

where the tensor *U_qkv_* ∈ ℝ *^D×3D^* projects each spot representation *r^i^* to three *D* dimensional vectors, *q^i^* (query), *k^i^* (key), and *v^i^* (value). The dot product between every query and all the keys, scaled by 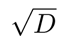, passes through a softmax function to obtain attention scores, *A*. The attention scores are the weighting of neighbor spots, which are then multiplied by their values, *v*, and each spot obtains a weighed representation with *D* dimensions. The second block further transforms the representation using a two-layer fully connected network (MLP) with 128 and 64 nodes. GELU [4] activation function is used to introduce non-linearity. Dropout [5] with rate 0.1 is applied after each layer.

Layer normalization (LN) [6] is applied before every block, and residual connection [7] after every block. Taking together, one encoder layer can be described as:

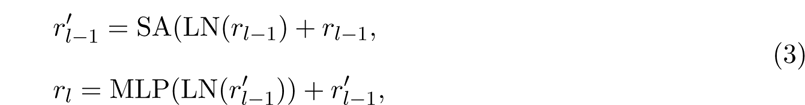

each encoder layer uses the output from the previous layer, *r_l-_*_1_, and generates the next, *r_l_*. The representation of the center spot from the last encoder layer, 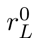, is taken as the input of the classifier.

### Classifier

We use the classifier to predict for a spot the gradient direction from it to its cell center and whether it is within a cell or outside a cell. The classifier has three components: (*i*) The spot representation 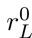 is first transformed using a two-layer MLP with 1024 and 256 nodes. LN is applied before the transformation. The output from the MLP is then used as input to two fully connected layers. (*ii*) One layer connects to a softmax function that outputs the probabilities, *d̂*, for each of the 16 possible directions to the cell center. (*iii*) Another layer generates a scalar output, *ŷ*, which is the object probability of the spot. Multi-class cross entropy is used as the loss function for the direction output:

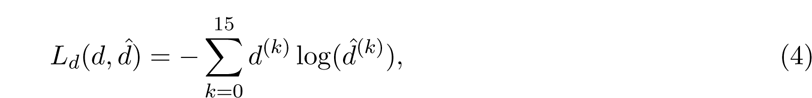

where *d* is the one-hot encoded direction label, where the bit with 1 indicates the correct direction. The binary cross entropy loss is used for the object probability:

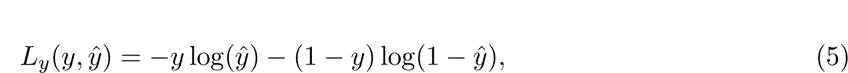

where *y* is a binary label indicating whether the spot is part of a cell. Considering all the *M* spots in the training data, the overall loss function is:

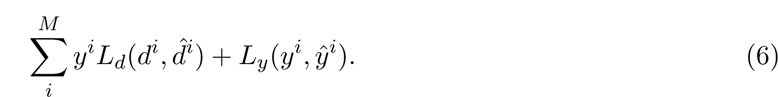

The direction loss is masked by the object label so the background spots will not contribute to this loss.

The selection of different model parts is discussed in Supplementary Figure 12.

### Training data preparation for the deep model

Spots within cell nucleus regions and spots sampled from highly confident background regions are used for model training. All spots within nucleus masks are labeled as 1 for the object label. For the direction label, the center of each nucleus is first computed by averaging the X and Y coordinates of spots assigned to this nucleus from the staining image. The direction from each spot within the nucleus to the nucleus center, *d*, is then computed as follows:

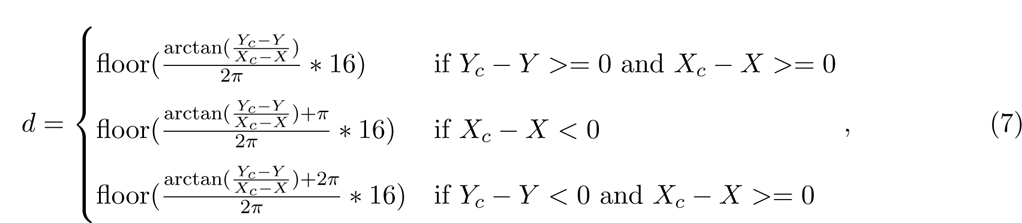

where *X* and *Y* are the coordinates of the spot for two axes, and *X_c_* and *Y_c_* are the coordinates of the nucleus center. The 16 direction classes evenly divide a full circle. On the other hand, the same number of spots as nucleus spots are sampled from background and labeled as 0 for the object label. The background spots have to meet the following two criteria: (*i*) The staining signal intensity of the corresponding pixel is smaller than a predefined threshold. (*ii*) The distance from the spot to any of the nucleus centers is greater than a predefined threshold.

### Cell boundary generation

Spot-level predictions are adjusted based on the locations of identified nuclei (Supplementary Note 11) and then smoothed (Supplementary Note 12), which are next used to group spots to cells. Spots with object probabilities smaller than 0.1 are determined to be background. The direction vectors of the rest of spots are treated as gradients and the gradient flow tracking algorithm [8] is performed to segment cells. In the algorithm, the vectors flow toward a sink, which corresponds to the center of the nuclei for each cell. Starting from a spot *b* = (*X, Y*), the next spot *b’* to which the flow is directed is selected using:

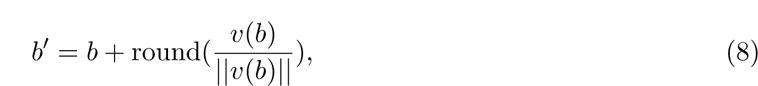

where *v*(*b*) is the gradient vector at *b*. When the angle between two consecutive steps is equal to or greater than 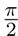, the gradient flow tracking procedure stops, and a sink is reached. The angle is computed as:

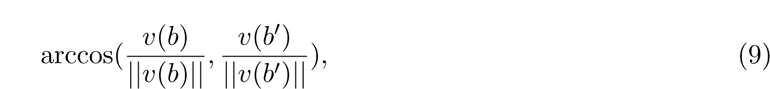

The set of spots that flow to the same sink produces an attraction basin of the sink. If the euclidean distance between two sinks is less than 3.5 spots, the attraction basins of the two sinks are combined together to obtain a larger attraction basin. An attraction basin with at least a certain number of spots is segmented as a cell (Supplementary Note 13).

### Correlation benchmark

We evaluate the performance of different methods by examining the transcriptional correlation between different regions of a segmentation mask. Specifically, for each nucleus detected using the staining, a target cell *c* is assigned using our cell segmentation by selecting the cell with the largest overlap with the source nucleus. A similar approach is used to select *c^0^* based on the cell segmentation of compared method. Gene expression profiles considering all the genes for three regions are computed: (*i*) intersection between *c* and *c*’, denoted as *x_int_*, (*ii*) region covered by *c* but not *c*’, denoted as *x_c_*, (*iii*) and region covered by *c*’ but not *c*, denoted as 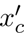. Then the Pearson linear correlation is used to measure the similarity of gene expression between *x_c_* and *x_int_*, as well as 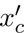 and *x_int_*. The segmentation is said to be better if its difference region has a higher correlation with the intersection region.

### Sub-cellular analysis

To perform sub-cellular analysis in SCS cell segmentation, we divided each cell mask into two regions, a nucleus region and a cytoplasm region. The nucleus region was identified using Watershed from the staining image data, while the cytoplasm region is the rest of the cell mask identified by SCS. RNAs detected in all spots within each region were aggregated to generate a gene expression profile, which was then summarized in a region by gene matrix. Genes whose RNAs localize differentially between two groups of regions were identified using t-test from the matrix.

For the Stereo-seq dataset, genes with *P*-values *<* 0.01 and fold changes greater than 1.3 were identified from each group. We then searched them in the RNALocate v2.0 database [9] for experimental evidence of RNA localization. As fewer genes were identified when using the Seq-scope dataset, likely due to smaller number of cells and challenges involved in identification of nucleus regions from H&E images, the top 100 genes with the smallest *P*-values were identified from each group and searched for experimental evidence.

### Application to Stereo-seq

#### Stereo-seq sequencing section

The Stereo-seq dataset captures a whole adult mouse brain slice in a single section (an area of around 5.3 mm×7.0 mm). The barcoded spots are arranged in a grid with a distance of 0.5 *µ*m between spots. Due to the large size of the assay, we cut it into small patches for model training, each with an area of 600 *µ*m×600 *µ*m (1200 spots×1200 spots) resulting in 87 patches, and processed one patch at a time. To obtain stable gene compositions for spots, we merged 3×3 spots into one spot. Therefore, each merged spot aggregates RNAs detected in a 1.5 *µ*m×1.5 *µ*m region. We computed the 2,000 variable genes across the spots using Scanpy [10], which decide the scope of genes in the expression profiles of spots.

#### Stereo-seq staining

The mouse brain tissue section was imaged with nucleic acid staining that visualized regions of nuclei. The stained image was aligned with the RNA assay in the original paper and we further refined the alignment using the linear transformation.

### Application to Seq-scope

#### Seq-scope sequencing sections

The Seq-scope dataset contains four tissue sections from mouse liver. Each section captures around a 0.93 mm×0.80 mm region of the tissue. The center-to-center distance of barcoded spots is similar to the distance for the Stereo-seq data, 0.5 *µ*m on average. Similar to Stereo-seq, we merged spots in each 1.5 *µ*m×1.5 *µ*m region into one spot. We again computed the 2,000 variable genes across the spots for each section.

#### Stereo-seq staining

The hematoxylin and eosin (H&E) image is provided for each of the sequencing sections. Therefore, the entire cell bodies are visualized in the imaging data. The stained image has been manually aligned with the RNA assay in the Stereo-seq paper. As a result, no further transformation was performed. We converted the H&E images to one channel grayscale images for the processing by Watershed.

## Data availability

All data used in this study have been previously published. The spatial transcriptomics data and nucleus staining images for the Stereo-seq dataset are available in the MOSTA data portal (https://db.cngb.org/stomics/mosta/download/) with file names Mouse brain Adult GEM bin1.tsv.gz and Mouse brain Adult.tif. The spatial transcriptomics data for the Seq-scope dataset are available in the Gene Expression Omnibus (GEO) database under accession number GSE169706. Tiles 2104, 2105, 2106, and 2107 were used in this study. The H&E staining images for the Seq-scope dataset are available at Deep Blue Data (https://doi.org/10.7302/cjfe-wa35). The seq-FISH+ NIH/3T3 cell line data are available at Zenodo (https://doi.org/10.5281/zenodo.2669683). The spatial transcriptomics data of the MERFISH human brain dataset are available at Dryad (https://datadryad.org/stash/dataset/doi:10.5061/dryad.x3ffbg7mw). The section of H22.26.401.MTG.4000 was used in this study. Experimental RNA subcellular localization data are available in the RNALocate v2.0 database (https://www.rna-society.org/rnalocate/download.html).

## Code availability

The source code of SCS is public available at https://github.com/chenhcs/SCS. The following open source Python (3.9.7) packages were used to build SCS: anndata (0.7.5), matplotlib (3.5.0), numpy (1.22.4), pandas (1.3.4), scanpy (1.8.2), scikit-learn (1.0.1), scipy (1.7.2), tensorflow (2.8.2). The open source software Spateo (0.0.0) was used to align staining image pixels with spatial transcriptomics spots. Watershed implemented in Spateo (0.0.0), open source Python packages DeepCell (0.12.3), Cellpose (2.1.1), StarDist (0.8.3), Baysor (0.5.2), and JSTA (0.0.0) were applied to segment staining images and compared with SCS.

---

**Extended Data Figure 1.**
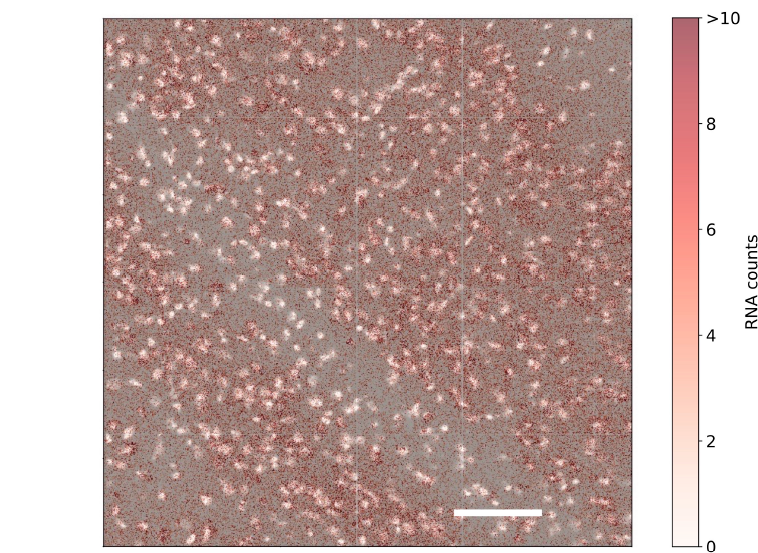

**Extended Data Figure 2.**
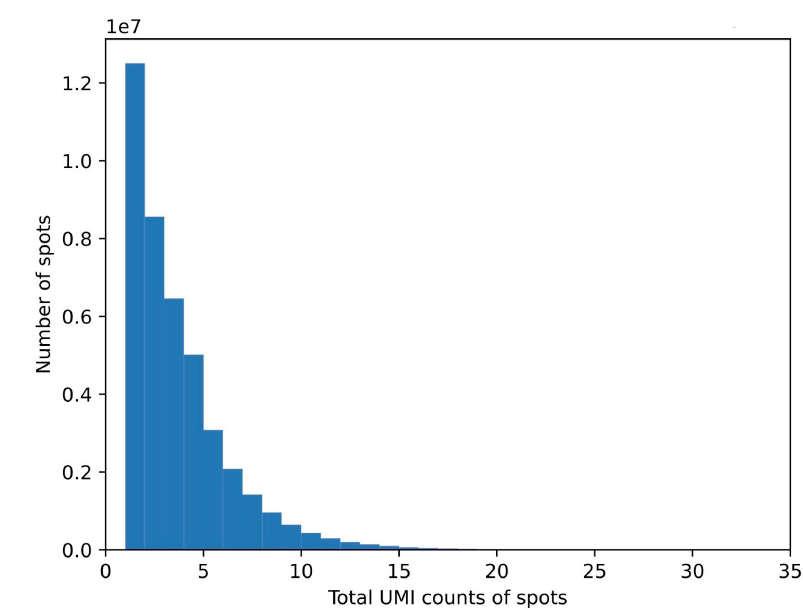

**Extended Data Figure 3.**
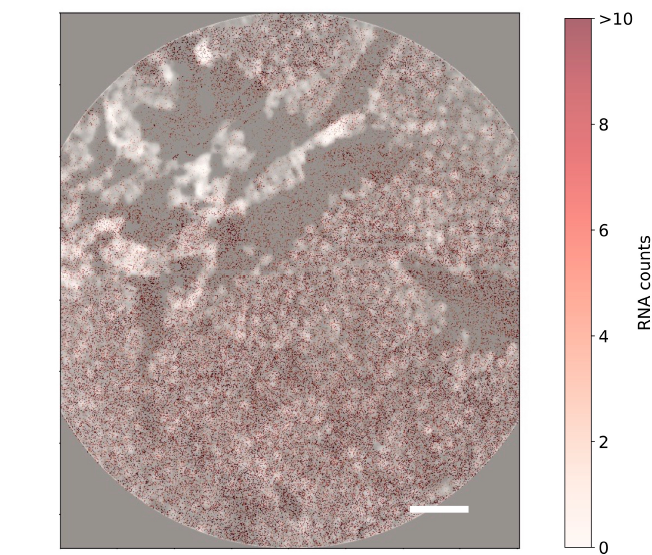

**Extended Data Figure 4.**
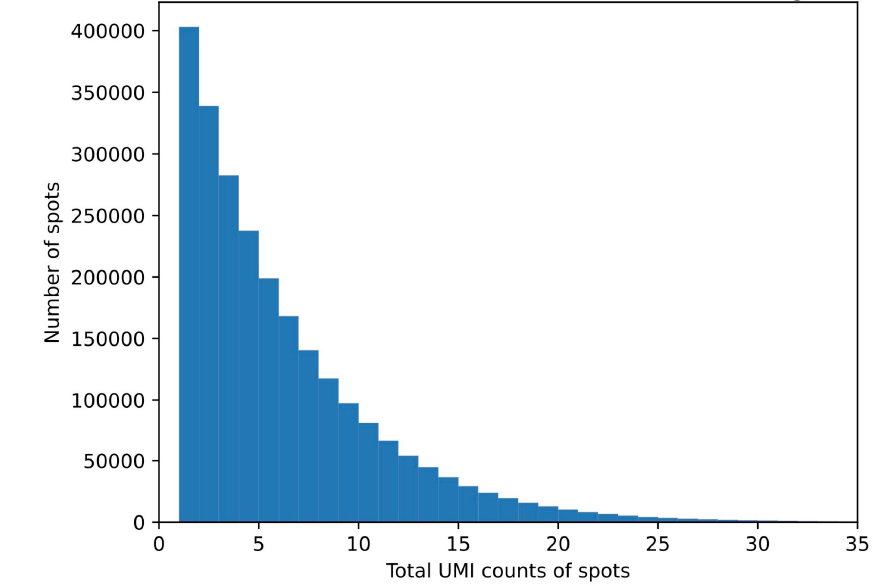

**Extended Data Figure 5.**
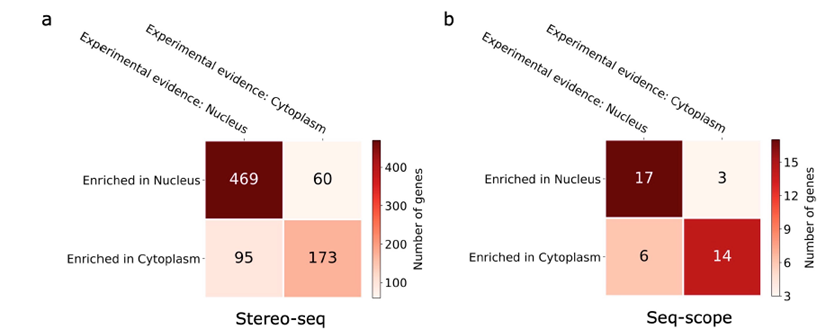

## References

[1] Dongshunyi Li, Jun Ding, and Ziv Bar-Joseph. Identifying signaling genes in spatial single-cell expression data. Bioinformatics, 37(7):968–975, 2021.

[2] Haotian Teng, Ye Yuan, and Ziv Bar-Joseph. Clustering spatial transcriptomics data. Bioin-formatics, 38(4):997–1004, 2022.

[3] Patrik L Ståhl, Fredrik Salmén, Sanja Vickovic, Anna Lundmark, José Fernández Navarro, Jens Magnusson, Stefania Giacomello, Michaela Asp, Jakub O Westholm, Mikael Huss, et al. Visualization and analysis of gene expression in tissue sections by spatial transcriptomics. Science, 353(6294):78–82, 2016.

[4] Ao Chen, Sha Liao, Mengnan Cheng, Kailong Ma, Liang Wu, Yiwei Lai, Xiaojie Qiu, Jin Yang, Jiangshan Xu, Shijie Hao, et al. Spatiotemporal transcriptomic atlas of mouse organogenesis using dna nanoball-patterned arrays. Cell, 185(10):1777–1792, 2022.

[5] Chun-Seok Cho, Jingyue Xi, Yichen Si, Sung-Rye Park, Jer-En Hsu, Myungjin Kim, Goo Jun, Hyun Min Kang, and Jun Hee Lee. Microscopic examination of spatial transcriptome using seq-scope. Cell, 184(13):3559–3572, 2021.

[6] Monica T Dayao, Maigan Brusko, Clive Wasserfall, and Ziv Bar-Joseph. Membrane marker selection for segmenting single cell spatial proteomics data. Nature communications, 13(1):1–10, 2022.

[7] Viktor Petukhov, Rosalind J Xu, Ruslan A Soldatov, Paolo Cadinu, Konstantin Khodosevich, Jeffrey R Moffitt, and Peter V Kharchenko. Cell segmentation in imaging-based spatial transcriptomics. Nature Biotechnology, 40(3):345–354, 2022.

[8] Russell Littman, Zachary Hemminger, Robert Foreman, Douglas Arneson, Guanglin Zhang, Fernando Gómez-Pinilla, Xia Yang, and Roy Wollman. Joint cell segmentation and cell type annotation for spatial transcriptomics. Molecular systems biology, 17(6):e10108, 2021.

[9] Serge Beucher. Use of watersheds in contour detection. In Proceedings of the International Workshop on Image Processing, pages 17–21. CCETT, 1979.

[10] Carsen Stringer, Tim Wang, Michalis Michaelos, and Marius Pachitariu. Cellpose: a generalist algorithm for cellular segmentation. Nature methods, 18(1):100–106, 2021.

[11] Dylan Bannon, Erick Moen, Morgan Schwartz, Enrico Borba, Takamasa Kudo, Noah Greenwald, Vibha Vijayakumar, Brian Chang, Edward Pao, Erik Osterman, et al. Deepcell kiosk: scaling deep learning–enabled cellular image analysis with kubernetes. Nature methods, 18(1):43–45, 2021.

12. Uwe Schmidt, Martin Weigert, Coleman Broaddus, and Gene Myers. Cell detection with star-convex polygons. In International Conference on Medical Image Computing and Computer-Assisted Intervention, pages 265–273. Springer, 2018.

[13] Noah F Greenwald, Geneva Miller, Erick Moen, Alex Kong, Adam Kagel, Thomas Dougherty, Christine Camacho Fullaway, Brianna J McIntosh, Ke Xuan Leow, Morgan Sarah Schwartz, et al. Whole-cell segmentation of tissue images with human-level performance using large-scale data annotation and deep learning. Nature biotechnology, 40(4):555–565, 2022.

[14] Joshua P Gilman, Maria Medalla, and Jennifer I Luebke. Area-specific features of pyramidal neurons—a comparative study in mouse and rhesus monkey. Cerebral Cortex, 27(3):2078–2094, 2017.

[15] Leland McInnes, John Healy, and James Melville. Umap: Uniform manifold approximation and projection for dimension reduction. arXiv preprint arXiv:1802.03426, 2018.

[16] Khan L Junatas, Zbyněk Tonar, Tereza Kubíková, Václav Liška, Richard Pálek, Patrik Mik, Milena Králíčková, and Kirsti Witter. Stereological analysis of size and density of hepatocytes in the porcine liver. Journal of Anatomy, 230(4):575–588, 2017.

[17] Chee-Huat Linus Eng, Michael Lawson, Qian Zhu, Ruben Dries, Noushin Koulena, Yodai Takei, Jina Yun, Christopher Cronin, Christoph Karp, Guo-Cheng Yuan, et al. Transcriptome-scale super-resolved imaging in tissues by rna seqfish+. Nature, 568(7751):235–239, 2019.

[18] Rongxin Fang, Chenglong Xia, Jennie L Close, Meng Zhang, Jiang He, Zhengkai Huang, Aaron R Halpern, Brian Long, Jeremy A Miller, Ed S Lein, et al. Conservation and divergence of cortical cell organization in human and mouse revealed by merfish. Science, 377(6601):56–62, 2022.

[19] Dylan M Cable, Evan Murray, Vignesh Shanmugam, Simon Zhang, Luli S Zou, Michael Diao, Haiqi Chen, Evan Z Macosko, Rafael A Irizarry, and Fei Chen. Cell type-specific inference of differential expression in spatial transcriptomics. Nature methods, 19(9):1076–1087, 2022.

[20] Ye Yuan and Ziv Bar-Joseph. Gcng: graph convolutional networks for inferring gene interaction from spatial transcriptomics data. Genome biology, 21(1):1–16, 2020.

[21] Yue Qin, Edward L Huttlin, Casper F Winsnes, Maya L Gosztyla, Ludivine Wacheul, Marcus R Kelly, Steven M Blue, Fan Zheng, Michael Chen, Leah V Schaffer, et al. A multi-scale map of cell structure fusing protein images and interactions. Nature, 600(7889):536–542, 2021.

[22] Tianyu Cui, Yiying Dou, Puwen Tan, Zhen Ni, Tianyuan Liu, DuoLin Wang, Yan Huang, Kaican Cai, Xiaoyang Zhao, Dong Xu, et al. Rnalocate v2. 0: an updated resource for rna subcellular localization with increased coverage and annotation. Nucleic acids research, 50(D1):D333–D339, 2022.

[23] Lisa Korostowski, Natalie Sedlak, and Nora Engel. The kcnq1ot1 long non-coding rna affects chromatin conformation and expression of kcnq1, but does not regulate its imprinting in the developing heart. PLOS Genetics, 8(9):e1002956, 2012.

[24] Christine M Clemson, John N Hutchinson, Sergio A Sara, Alexander W Ensminger, Archa H Fox, Andrew Chess, and Jeanne B Lawrence. An architectural role for a nuclear noncoding rna: Neat1 rna is essential for the structure of paraspeckles. Molecular cell, 33(6):717–726, 2009.

[25] Jan RT Van Weering, Ruud F Toonen, and Matthijs Verhage. The role of rab3a in secretory vesicle docking requires association/dissociation of guanidine phosphates and munc18-1. PLoS One, 2(7):e616, 2007.

[26] Natali L Chanaday and Ege T Kavalali. Synaptobrevin-2 dependent regulation of single synap-tic vesicle endocytosis. Molecular biology of the cell, 32(19):1818–1823, 2021.

[27] Krysta L Engel, Ankita Arora, Raeann Goering, Hei-Yong G Lo, and J Matthew Taliaferro. Mechanisms and consequences of subcellular rna localization across diverse cell types. Traffic, 21(6):404–418, 2020.

## References

[1] Ao Chen, Sha Liao, Mengnan Cheng, Kailong Ma, Liang Wu, Yiwei Lai, Xiaojie Qiu, Jin Yang, Jiangshan Xu, Shijie Hao, et al. Spatiotemporal transcriptomic atlas of mouse organogenesis using dna nanoball-patterned arrays. Cell, 185(10):1777–1792, 2022.

[2] Chun-Seok Cho, Jingyue Xi, Yichen Si, Sung-Rye Park, Jer-En Hsu, Myungjin Kim, Goo Jun, Hyun Min Kang, and Jun Hee Lee. Microscopic examination of spatial transcriptome using seq-scope. Cell, 184(13):3559–3572, 2021.

[3] Ashish Vaswani, Noam Shazeer, Niki Parmar, Jakob Uszkoreit, Llion Jones, Aidan N Gomez, Lukasz Kaiser, and Illia Polosukhin. Attention is all you need. Advances in neural information processing systems, 30, 2017.

[4] Dan Hendrycks and Kevin Gimpel. Gaussian error linear units (gelus). arXiv preprint arXiv:1606.08415, 2016.

[5] Nitish Srivastava, Geoffrey Hinton, Alex Krizhevsky, Ilya Sutskever, and Ruslan Salakhutdinov. Dropout: a simple way to prevent neural networks from overfitting. The journal of machine learning research, 15(1):1929–1958, 2014.

[6] Jimmy Lei Ba, Jamie Ryan Kiros, and Geoffrey E Hinton. Layer normalization. arXiv preprint arXiv:1607.06450, 2016.

[7] Kaiming He, Xiangyu Zhang, Shaoqing Ren, and Jian Sun. Deep residual learning for image recognition. In Proceedings of the IEEE conference on computer vision and pattern recognition, pages 770–778, 2016.

[8] Gang Li, Tianming Liu, Ashley Tarokh, Jingxin Nie, Lei Guo, Andrew Mara, Scott Holley, and Stephen TC Wong. 3d cell nuclei segmentation based on gradient flow tracking. BMC cell biology, 8(1):1–10, 2007.

[9] Tianyu Cui, Yiying Dou, Puwen Tan, Zhen Ni, Tianyuan Liu, DuoLin Wang, Yan Huang, Kaican Cai, Xiaoyang Zhao, Dong Xu, et al. Rnalocate v2. 0: an updated resource for rna subcellular localization with increased coverage and annotation. Nucleic acids research, 50(D1):D333–D339, 2022.

[10] F Alexander Wolf, Philipp Angerer, and Fabian J Theis. Scanpy: large-scale single-cell gene expression data analysis. Genome biology, 19(1):1–5, 2018.

