## Supplementary Notes and Figures for "SCS: cell segmentation for high-resolution spatial transcriptomics"

##### Supplementary Note 1 Watershed segmentation with Spateo

The Watershed algorithm implemented in Spateo (https://spateo-release.readthedocs.io) [1] was used to segment nuclei as well as whole cell bodies from staining images. The algorithm first uses a combination of global and local thresholding to obtain a mask of objects (nuclei or cells). Then peaks are identified from the mask based on the intensity gradient defined on the image, which are used as Watershed markers. Masks of nuclei or cells are expanded from the markers. The cell segmentation uses a lower background threshold therefore expands the nuclei masks to cytoplasm regions. The Watershed cell segmentation is compared with SCS cell masks.

##### Supplementary Note 2 Cell segmentation with the deep learning based methods

Due to the lack of manual cell segmentation labels, we selected appropriate pretrained models of the deep learning based methods for performance evaluation. For StartDist [2], the model from its original publication was used, which was trained on the DSB 2018 nuclei segmentation challenge dataset. The cytoplasm model of Cellpose [3] was used for evaluation, while the nuclear model of DeepCell [4] was used, as the tests of other models of these two methods on the two datasets showed that they failed to identify most of the cells. The Mesmer [5] model integrated in DeepCell was also used for comparison (Supplementary Figs. 11-13).

##### Supplementary Note 3 Cell filtering for the correlation benchmark

When evaluating the accuracy of two cell segmentation using the correlation benchmark, a low correlation can result from one region having only a few detected RNAs aggregated. For example, one cell mask is almost contained in another cell mask, the difference region for the first cell mask will be very small. To make fair comparisons, we considered only cell mask pairs that all the three regions (the intersection region and two difference regions) have at least 100 RNA molecules aggregated.

#### Supplementary Note 4 Calculation of the cell diameter

We assume that the shape of each cell in the tissue section is round, and then calculate the diameter of each cell through the area of its segmented cell mask (number of spots covered).

#### Supplementary Note 5 Defining novel cell predictions

We define novel cell predictions of method $A$ when compared with method $B$ as: ($i$) Cells identified by method $A$ that do not overlap with any cell identified by method $B$, ($ii$) Cells identified by method $A$ that are merged by a cell from method $B$. Only cells that have 90% area covered by another cell from the other method are considered as being merged by the cell. The cells being merged except for the largest one are considered novel predictions of method $A$.

#### Supplementary Note 6 Cell type annotation

We transferred cell type labels from the Mouse Steady State dataset in Liver Cell Atlas [6] to the SCS segmented cells on the Seq-scope dataset. The intersection of genes from two datasets was taken to make them defined on the same variables. The ingest data integration function from the Scanpy package [7] was then used for label transferring and UMAP projection of cells, in which the Mouse Steady State dataset was treated as reference data and the segmented cells on the Seq-scope dataset were used as new data. The function fits a model of PCA combined with a neighbor lookup search tree on the reference data and uses it to project new data.

The cell type labels from an annotated scRNA-seq dataset of adult mouse brain [8] were transferred to the SCS segmented cells on the Stereo-seq dataset. The same integration method as Seq-scope was used for the Stereo-seq dataset.

#### Supplementary Note 7 Evaluation on the seqFISH+ dataset

The seqFISH+ dataset for NIH/3T3 cells [9] contains 17 tiles. Each tile sizes about 200 $\mu$m $\times$ 200 $\mu$m, with 10,000 genes profiled in each tile. This dataset also has paired DAPI nuclei staining images for each profiled tile. To transform the detected RNA points to grid-shaped spots that SCS can process, we aggregated RNAs detected in each 0.4 $\mu$m $\times$ 0.4 $\mu$m region into a spot, and then run SCS on the transformed data. We again computed the 2,000 variable genes across the spots for each tile, which decide the scope of genes in the expression profiles of spots. For this dataset, manually annotated cell segmentation ground-truth is available from the original publication [9]. Thus to evaluate the performance of different methods on this data, we quantified the agreement of cell segmentations with the ground truth (Intersection over Union (IoU)).

#### Supplementary Note 8 Evaluation on the MERFISH dataset

The MERFISH dataset for the human brain tissue was collected from [10]. This dataset profiled 4,000 genes for a human brain tissue section of size around 4816 $\mu$m $\times$ 4480 $\mu$m and was paired with DAPI nuclei staining images. We again transformed the data into grid-shaped spots, each covering a 1.68 $\mu$m $\times$ 1.68 $\mu$m region. Similar to the Stereo-seq dataset, we cut the MERFISH section into small patches for model training, each with an area of 672 $\mu$m$\times$672 $\mu$m (1200 spots$\times$1200 spots), and processed one patch at a time. The 2,000 variable genes across the spots for each patch were computed and used for the expression profiles of spots. As no ground-truth segmentation is available for this dataset, we used the correlation metric for performance evaluation.

#### Supplementary Note 9 Cell segmentation with Baysor

Baysor was run on the same variable genes identified in SCS. For Stereo-seq and MERFISH, Baysor was run on the same patch splits as SCS. We set hyperparameters of Baysor as default. To make the method achieve its best performance, we allow Baysor to use the nucleus segmentation results to estimate the key hyperparameter "scale". Six patches in Stereo-seq and two patches in MERFISH were unfinished given they did not meet the minimum requirement to estimate the "scale" hyperparameter according to Baysor. Same as SCS, predicted cells of Baysor that cover less than 200 spots were ignored.

#### Supplementary Note 10 Cell segmentation with JSTA

JSTA requires both the input of image based segmentation priors and single cell RNA-seq (scRNA-seq) dataset of the same tissue with cell type annotations. For the image based segmentation priors, we provide JSTA with the Watershed nucleus segmentations generated from the first step of SCS. For the scRNA-seq dataset, we collected single cell data for the same tissue of the spatial transcriptomics data. Specifically, for the Stereo-seq dataset, we used an annotated scRNA-seq dataset of adult mouse brain [8]. We used a subsample of 50,650 cells selected using stratified sampling based on cell type labels. For the Seq-scope dataset, we used the annotated scRNA-seq dataset of mouse from the Liver Cell Atlas [6]. We used a subsample of 50,191 cells selected using stratified sampling based on cell type labels. For the MERFISH dataset, we used the SMART-seq dataset (https://portal.brain-map.org/atlases-and-data/rnaseq/human-mtg-smart-seq) used in the original publication of the MERFISH paper. For the seqFISH+ dataset, we used the single cells of the NIT/3T3 cell line collected from 10x Genomics datasets (https://www.10xgenomics.com/resources/datasets/293-t-and-3-t-3-cell-mixture-1-standard-1-1-0). Running a whole Seq-scope tile resulted in too much memory consumption in JSTA, so we cut a tile into four patches (upper left, upper right, lower left, and lower right) and ran each patch separately. For Stereo-seq and MERFISH, Baysor was run on the same patch splits as SCS.

We set hyparameters the same as in the tutorial notebook in JSTA GitHub repository except for *pixel_length* and *num_genes_to_keep.* To avoid memory leak and to be able to finish JSTA within reasonable amount of time, we set these two hyperparameters differently for different datasets, according to the instructions from JSTA. Specifically, For Seq-scope, we have *pixel_length*=3 and *num_genes_to_keep*=100. For other datasets, we have *pixel_length*=1 and *num_genes_to_keep*=200. One patch of the seqFISH+ dataset and sixteen patches of Stereo-seq dataset were unfinished due to errors in learning the cell type classifier in JSTA. Same as SCS, predicted cells of JSTA that cover less than 200 spots were ignored.

#### Supplementary Note 11 Spot-level prediction prior adjustment

Given the nuclei positions around a spot, some gradient directions become more likely to be predicted. We therefore correct direction predictions by adjusting the prior probabilities of direction classes [11] based on coordinates of surrounding nuclei and prior knowledge of cell sizes. Information are required the expected cell radius, $\mu_{size}$, and standard deviation of cell size, $\sigma_{size}$, with $\sigma_{size} = \mu_{size}/4$ by default. We use 7.5 $\mu$m as the expected cell radius for the Stereo-seq data on mouse brain cells and 10.0 $\mu$m as the expected cell radius for the Seq-scope data on mouse liver cells, according to previous findings in the literature [12, 13]. Without knowing coordinates of surrounding nuclei, the model assumes the prior probabilities of $y_d$ are uniformly distributed:

$$P_{old}(y_d = k) = \frac{1}{16}, \text{for } k \in [0, 15]. \tag{1}$$

However, given the coordinates of surrounding nuclei of a spot, choices of predictions are narrowed down to the directions where nuclei centers can be found in a certain range, specifically:

$$P_{new}(y_d = k) = \begin{cases} 0 & \text{if no nucleus within } \mu_{size} + \sigma_{size} \text{ on this direction} \\ \frac{1}{n} & \text{if nuclei within } \mu_{size} + \sigma_{size} \text{ on this direction} \end{cases}, \tag{2}$$

where $n$ is the number of directions that find nuclei. The predicted probability on direction $k$ for the spot is then corrected as:

$$P_{new}(y_d = k|x, s) = \frac{P_{old}(y_d = k|x, s)P_{new}(y_d = k)/P_{old}(y_d = k)}{\sum_{k'=0}^{15} P_{old}(y_d = k'|x, s)P_{new}(y_d = k')/P_{old}(y_d = k')}. \tag{3}$$

The direction class with the largest probability is taken that is then transformed back to the two dimensional direction vector.

#### Supplementary Note 12 Spot-level prediction smoothing

Noise presented in direction vectors or "holes" in the sequencing section (spots without any RNA detected) may lead to an unexpected termination of flow tracking. We therefore smooth the spot-level predictions by averaging the direction vectors and object probabilities of the eight immediate neighbors of each spot and the spot itself. The smoothing process is performed two rounds.

#### Supplementary Note 13 Removing small attraction basins

After performing the gradient flow tracking algorithm, attraction basins that cover less than 200 spots are ignored to avoid noise in the background from being identified as cells.

### Supplementary Figures

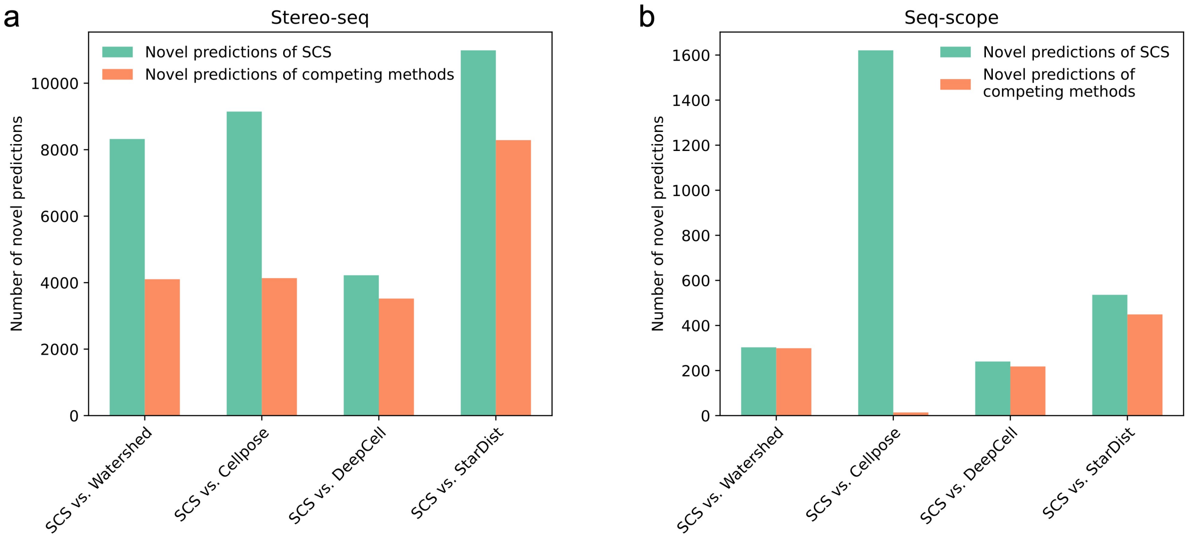

**Supplementary Figure 1: The number of novel predictions of different methods when compared against each other**. **a**, The comparison of the number of novel predictions between SCS and the other methods on the Stereo-seq dataset. **b**, The comparison on the Seq-scope dataset.

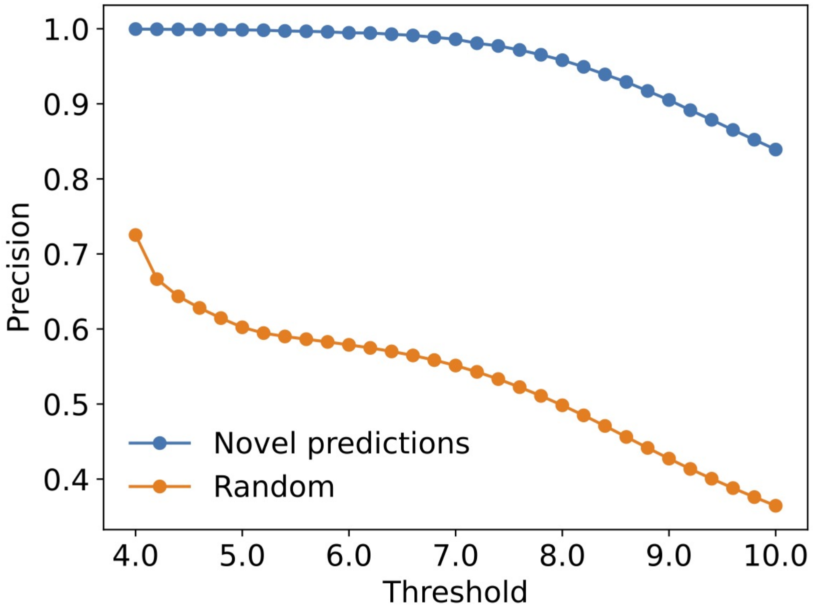

**Supplementary Figure 2: Precision of the novel cell predictions.** We estimated the precision of the novel cell predictions identified by SCS (cells that do not overlap with cells identified using Watershed segmentation) in the Stereo-seq dataset by using the intensity of the nucleus staining as a cutoff to select potential true positives. Specifically, using this method we consider cells whose average staining intensity is greater than a threshold as true positives and the rest as false positives. We tested thresholds ranging from the first quartile to the third quartile of the staining intensities of the background pixels (4.0 to 10.0), the precision of the novel SCS recovered cells ranges from 1.0 to 0.84, which is significantly higher than that of randomly picking a region from the background as a cell.

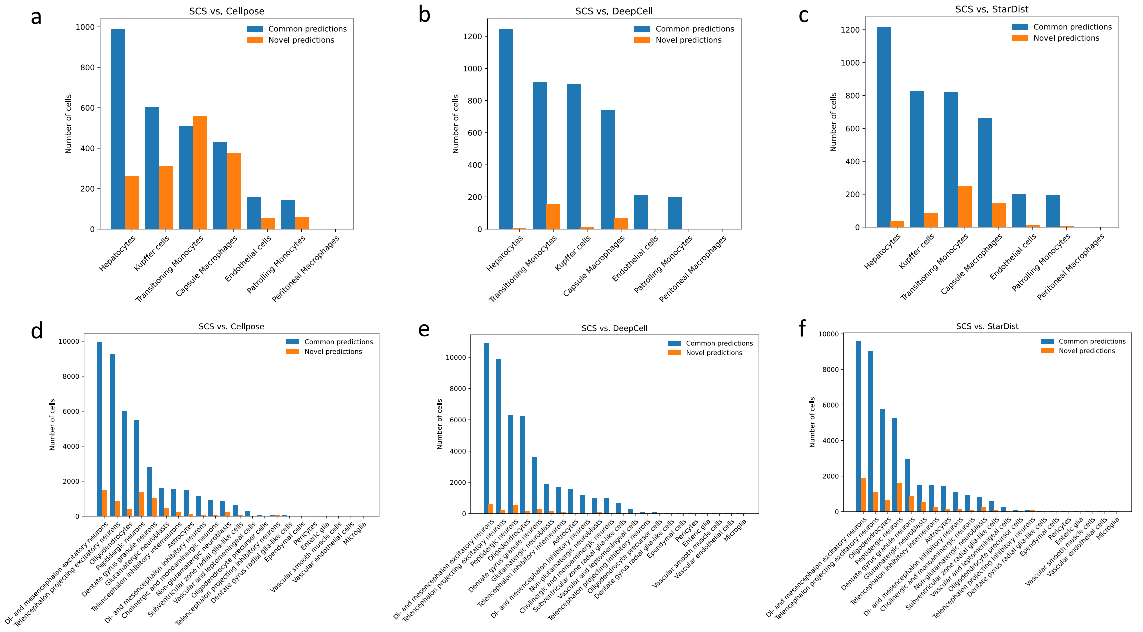

**Supplementary Figure 3: The distribution of novel cell predictions of SCS across different cell types.** **a**, The number of novel cell predictions by SCS compared to Cellpose vs. the number of cells that are commonly identified by SCS and Cellpose in different cell types for the Seq-scope dataset. **b**, The comparison for SCS vs. DeepCell on the Seq-scope dataset. **c**, The comparison for SCS vs. StarDist on the Seq-scope dataset. **d**, The comparison for SCS vs. Cellpose on the Stereo-seq dataset. **e**, The comparison for SCS vs. DeepCell on the Stereo-seq dataset. **f**, The comparison for SCS vs. StarDist on the Stereo-seq dataset.

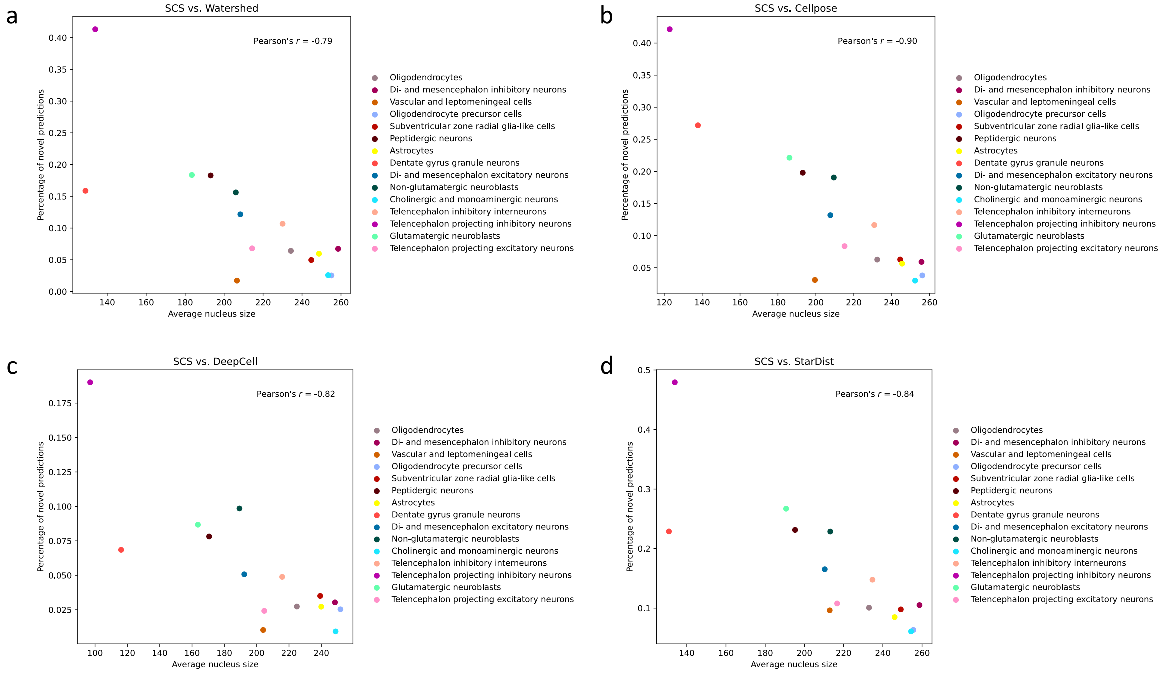

**Supplementary Figure 4: The correlation between the percentage of novel cells in different cell types with their average nucleus sizes on the Stereo-seq dataset**. **a**, Each dot shows for each cell type the percentage of novel cell predictions by SCS compared to Watershed and its average nucleus size. The two statistics show a negative correlation across cell types. **b**, The correlation analysis where the novel cell predictions are defined by comparing SCS segmented cells with Cellpose segmentation. **c**, The correlation analysis where the novel cell predictions are defined by comparing SCS segmented cells with DeepCell segmentation. **d**, The correlation analysis where the novel cell predictions are defined by comparing SCS segmented cells with StarDist segmentation. **a-d**, Only cell types with at least 50 cells are considered in this plot.

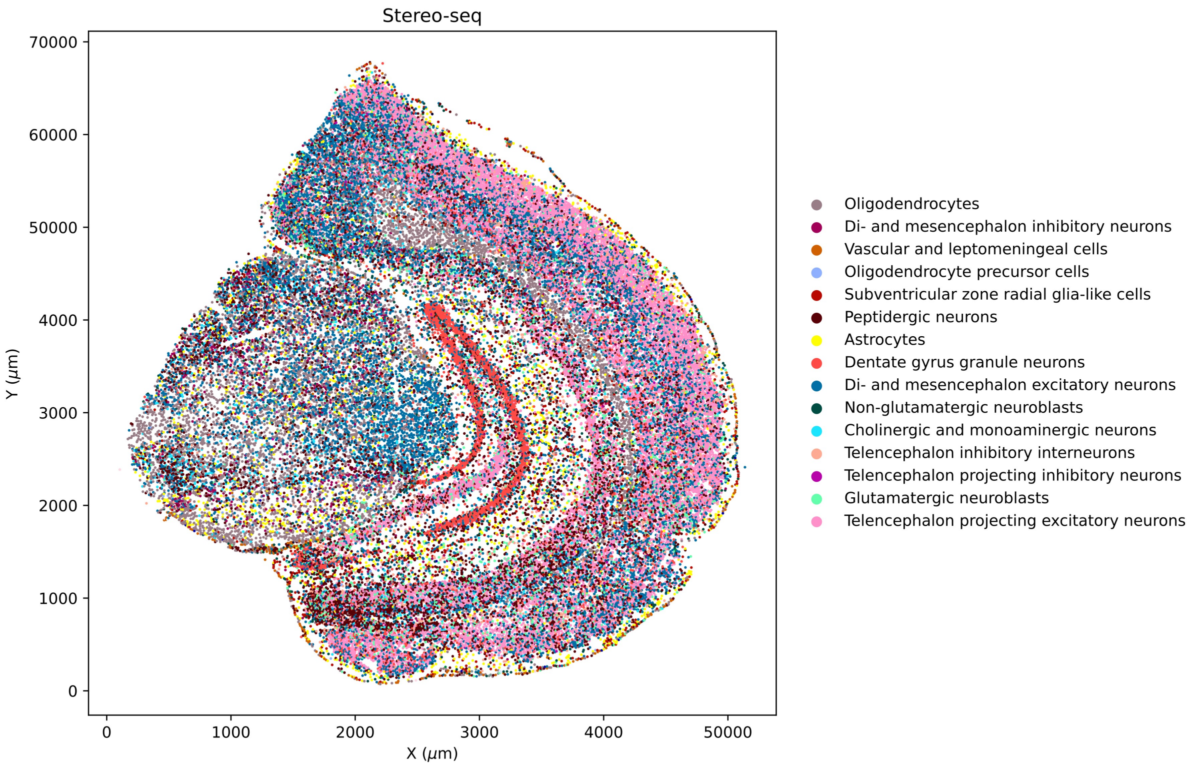

**Supplementary Figure 5: The spatial distribution of cell types of the SCS segmented cells in the Stero-seq dataset.** The cell type annotations are consistent with expected histological features. For example, the Dentate gyrus granule neurons mainly appear in the dentate gyrus region of the mouse brain.

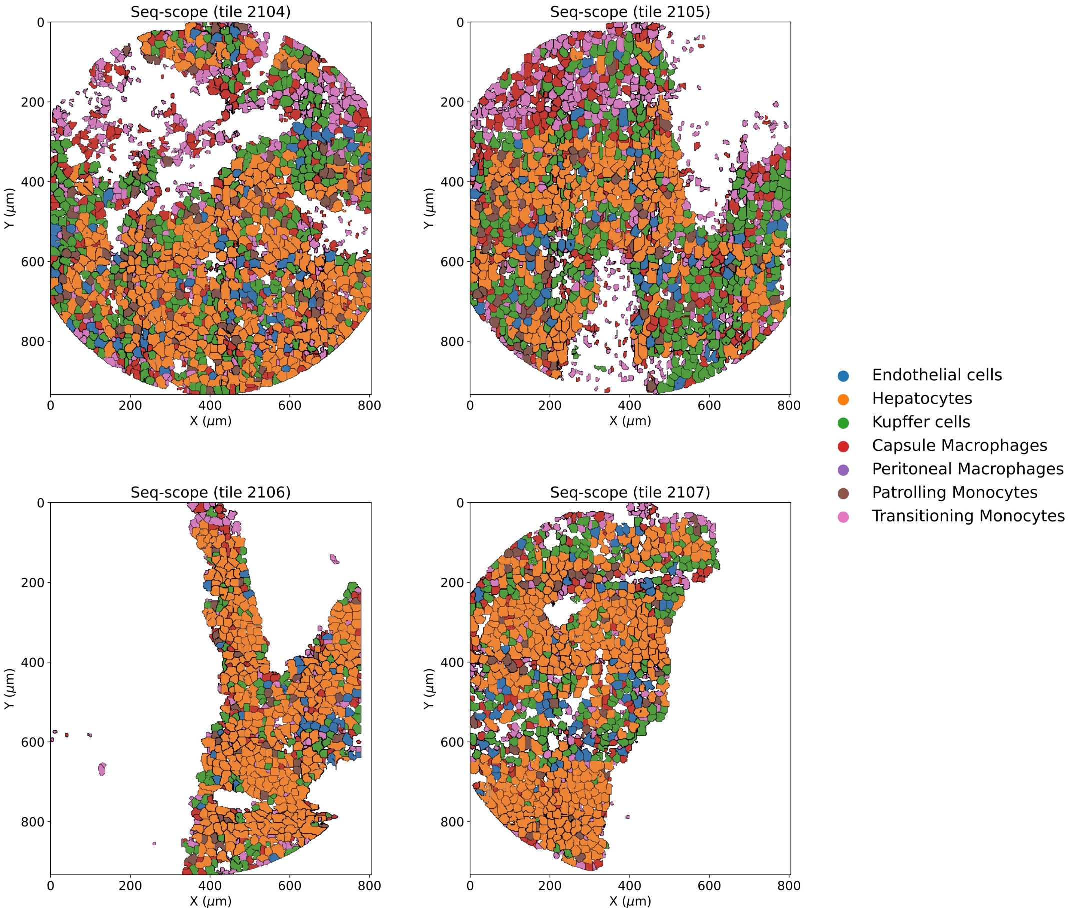

**Supplementary Figure 6:** The spatial distribution of cell types of the SCS segmented cells in the four tiles of the Seq-scope dataset.

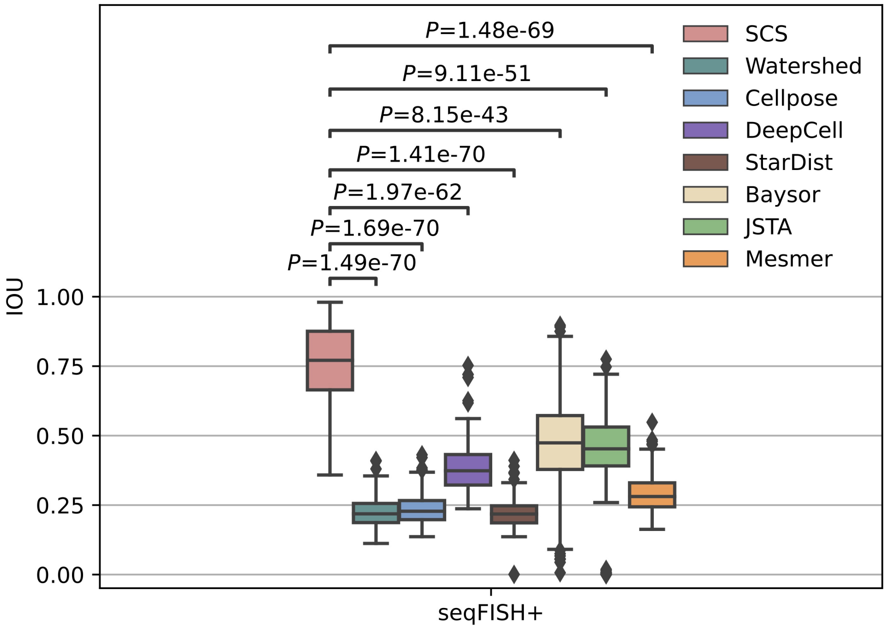

**Supplementary Figure 7: Evaluation on the seqFISH+ NIH/3T3 cell line dataset.** We quantified the agreement of cell segmentations with the ground truth (Intersection over Union (IoU)) to evaluate the performance of different methods on this dataset. The average IoU of SCS across all the cells in this dataset is 0.75, which is much higher than two previously proposed cell segmentation methods for spatial transcriptomics data Baysor (0.46) and JSTA (0.44), as well as all the other image based methods (Kruskal-Wallis tests; $N$=211 cells for SCS, $N$=211 cells for Watershed, $N$=211 cells for Cellpose, $N$=211 cells for DeepCell, $N$=211 cells for StarDist, $N$=211 cells for Baysor, $N$=200 cells for JSTA, one patch unfinished, $N$=211 cells for Mesmer). Boxplots show medians (horizontal line in each box), interquartile ranges (boxes), 1.5 interquartile (whiskers), and remaining point individuals.

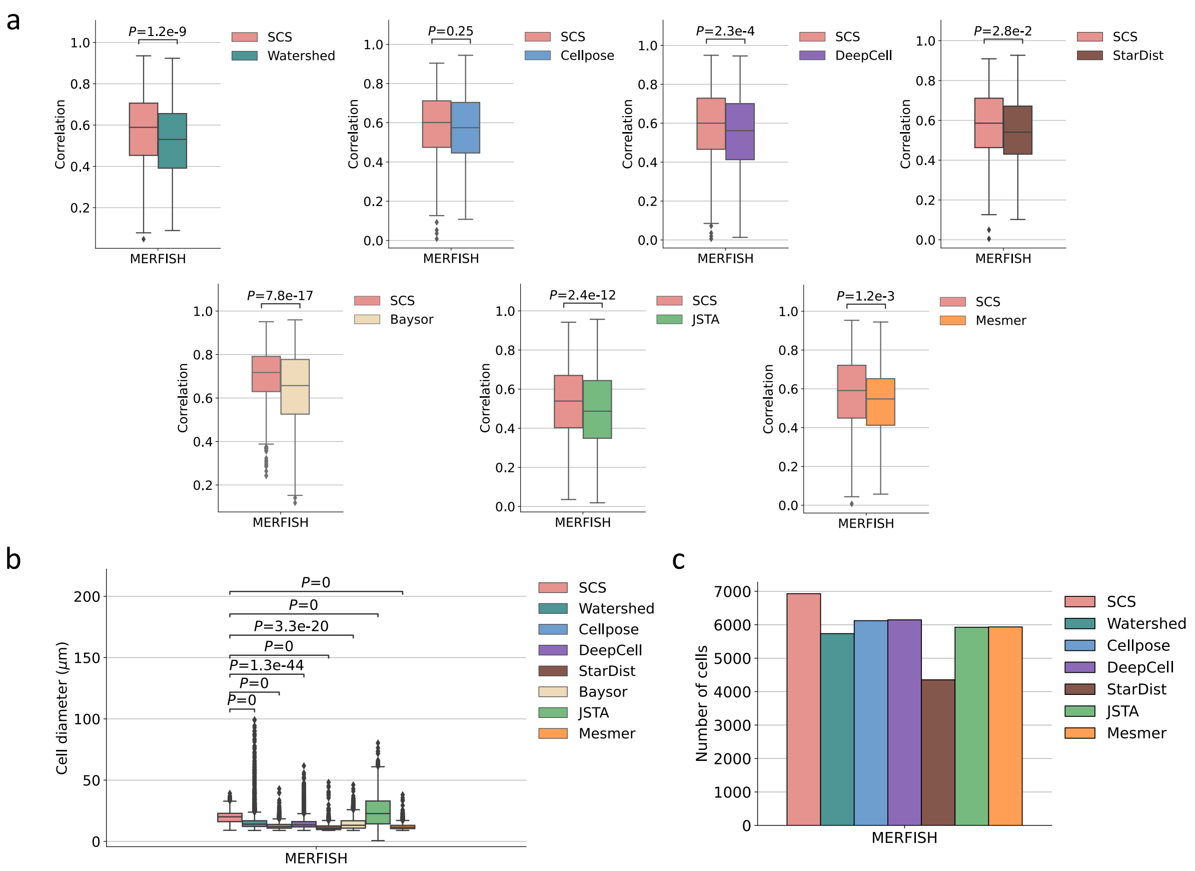

**Supplementary Figure 8: Evaluation on the MERFISH human brain dataset.** **a**, The comparisons between SCS and the other methods on the MERFISH dataset using the correlation benchmark. SCS achieved higher segmentation accuracy than all the other methods on this dataset (Wilcoxon signed-rank test, one-sided; $N$=800 cells for SCS vs. Watershed, $N$=434 cells for SCS vs. Cellpose, $N$=760 cells for SCS vs. DeepCell, $N$=234 cells for SCS vs. StarDist, $N$=973 cells for SCS vs. Baysor, $N$=1,787 cells for SCS vs. JSTA, $N$=335 cells for SCS vs. Mesmer). **b**, Comparison of the sizes of cells segmented by SCS and other methods (Kruskal-Wallis tests; $N$=6,930 cells for SCS, $N$=5,735 cells for Watershed, $N$=6,122 cells for Cellpose, $N$=6,147 cells for DeepCell, $N$=4,354 cells for StarDist, $N$=6,104 cells for Baysor, $N$=5,926 cells for JSTA, $N$=5,935 cells for Mesmer). As nucleus staining was used, the image based methods (Watershed, Cellpose, DeepCell, StarDist, and Mesmer) tend to segment only nucleus regions and thus underestimate cell sizes. **c**, The number of cells identified by different methods. As some patches are unfinished for Baysor (Supplementary Notes), its number of cells is not compared in the figure. **a-b**, Boxplots show medians (horizontal line in each box), interquartile ranges (boxes), 1.5 interquartile (whiskers), and remaining point individuals.

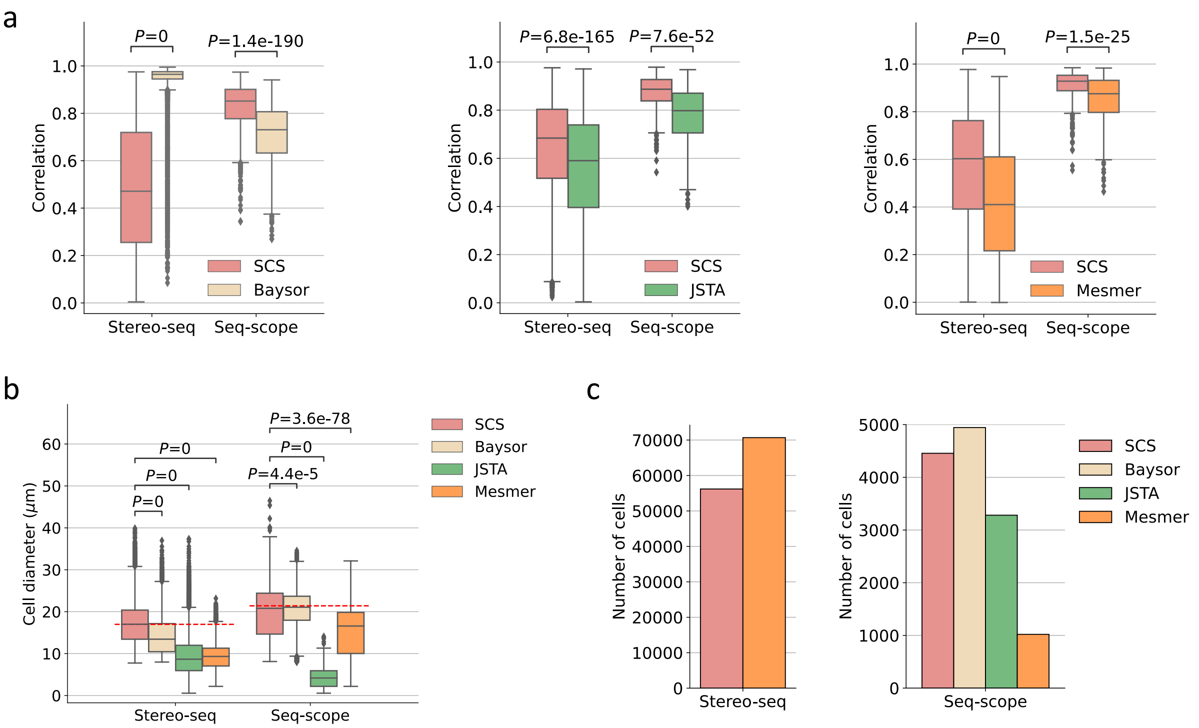

**Supplementary Figure 9: Additional comparisons with methods developed for spatial transcriptomics data, Baysor and JSTA, and another image-based model, Mesmer. a**, The comparisons between SCS and other methods on Stereo-seq and Seq-scope using the correlation benchmark. SCS significantly outperforms JSTA and Mesmer on both datasets and strongly outperforms Baysor on the Seq-scope dataset (Wilcoxon signed-rank tests, one-sided; for SCS vs. Baysor, $N$=13,098 cells on Stereo-seq, $N$=2,187 cells on Seq-scope; for SCS vs. JSTA, $N$=10,001 cells on Stereo-seq, $N$=636 cells on Seq-scope, for SCS vs. Mesmer, $N$=10,049 cells on Stereo-seq, $N$=536 cells on Seq-scope.). **b**, Comparison of the sizes of cells segmented by SCS and other methods (Kruskal-Wallis tests; for Stereo-seq, $N$=56,187 cells for SCS, $N$=38,295 cells for Baysor, $N$=41,925 for JSTA, $N$=70,679 cells for Mesmer; for Seq-scope, $N$=4,456 cells for SCS, $N$=4,944 cells for Baysor, $N$=3,281 cells for JSTA, $N$=1,020 cells for Mesmer). On the Stereo-seq dataset, Baysor achieves very high correlations likely because it only segmented nucleus regions. The red dashed lines show the expected cell diameters from the literature [12, 13]. **c**, The number of cells identified by different methods. For Stereo-seq, as some patches are unfinished for Baysor and JSTA (Supplementary Notes), their numbers of cells are not compared in the figure. **a-b**, Boxplots show medians (horizontal line in each box), interquartile ranges (boxes), 1.5 interquartile (whiskers), and remaining point individuals.

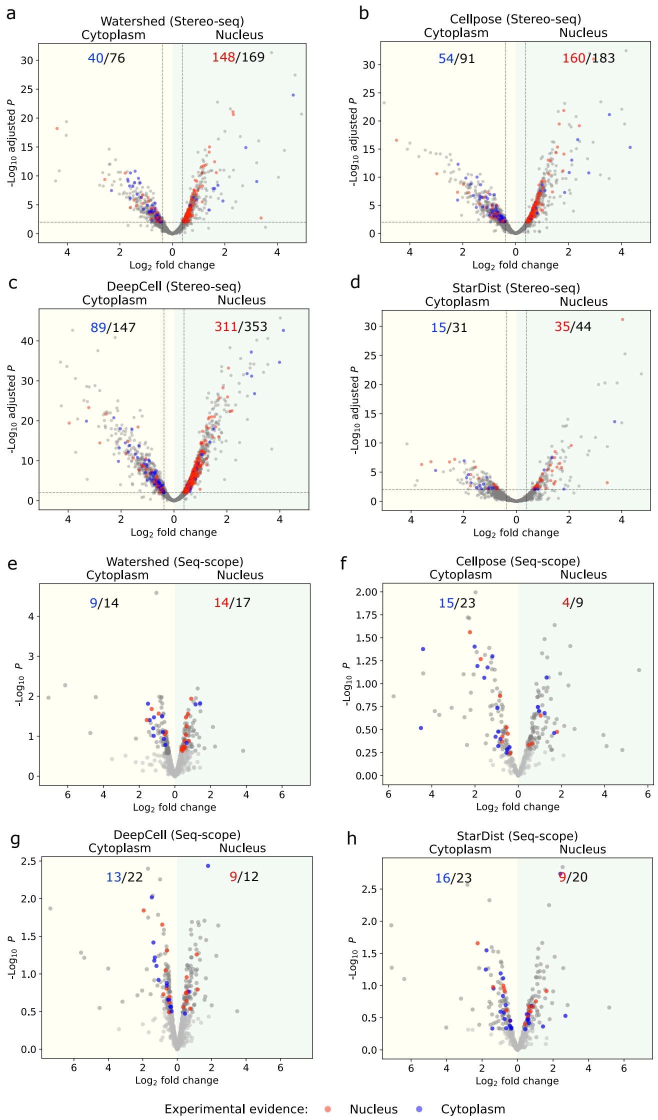

**Supplementary Figure 10: Subcellular analysis for the image based methods. a-d**, Volcano plots for the image based methods that show quantitative changes in expression levels for genes between the nucleus and cytoplasm regions of their segmented cells on the Stereo-seq dataset (t-tests, one-sided, the Benjamini-hochberg method was used to adjust $P$-values; Watershed: $N$=34,853 nucleus regions, $N$=35,792 cytoplasm regions, Cellpose: $N$=32,223 nucleus regions, $N$=34,563 cytoplasm regions, DeepCell: $N$=32,567 nucleus regions, $N$=39,197 cytoplasm regions, StarDist: $N$=31,717 nucleus regions, $N$=15,786 cytoplasm regions, for each method, subcellular regions with at least 100 genes were used for this analysis). **e-f**, Volcano plots for image based methods on the Seq-scope dataset (t-tests, one-sided, raw $P$-values are shown to avoid most of the $P$-values being corrected to the same value; Watershed: $N$=2, 038 nucleus regions, $N$=2,236 cytoplasm regions, Cellpose: $N$=1,710 nucleus regions, $N$=1,598 cytoplasm regions, DeepCell: $N$=2,573 nucleus regions, $N$=2,824 cytoplasm regions, StarDist: $N$=2,038 nucleus regions, $N$=2,236 cytoplasm regions, for each method, the subcellular regions with at least 100 genes are used for this analysis). **a-h**, The numbers on the left panel of each subfigure show the number of blue dots vs. the number of colored dots. The numbers on the right panel of each subfigure show the number of red dots vs. the number of colored dots.

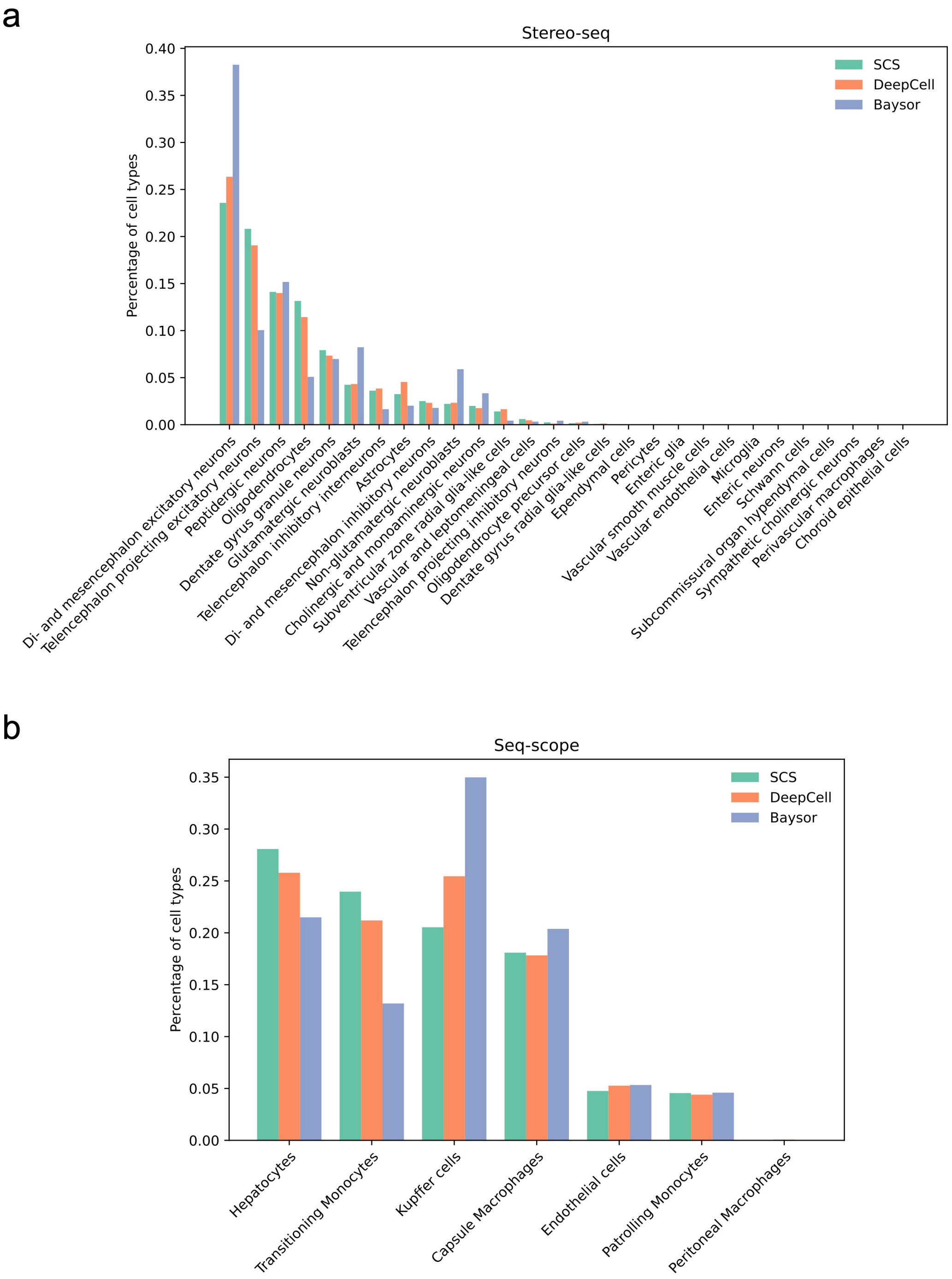

**Supplementary Figure 11: Cell type distributions of identified cells by different methods. a**, The comparison of cell type distributions for segmented cells identified by SCS, DeepCell, and Baysor on the Stereo-seq dataset. **b**, The comparison of cell type distributions for segmented cells identified by SCS, DeepCell, and Baysor on the Seq-scope dataset.

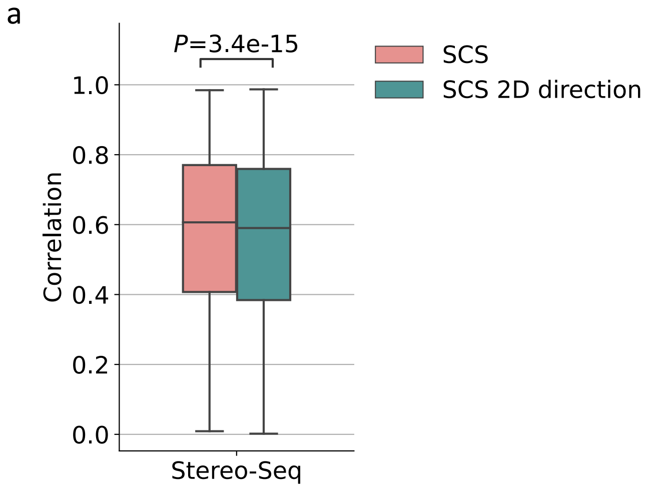

**Supplementary Figure 12: Testing of different options of model parts**. **a**, Comparison between different options for the direction representation of the model. The 16 dimensional categorical vector is replaced with a two-dimensional vector with the fixed length of 1 and real number in each bit. And the classification loss of the model is changed to regression loss (SCS 2D direction). We tested this model on the Stereo-seq dataset. Results indicate that the performance of the 16 dimensional categorical vector is significantly better than that of the two-dimensional vector (0.61 vs. 0.59, Wilcoxon signed-rank test, one-sided; $N$=32,928 cells). One possible explanation is that for the categorical (16 dimensions) representations, we can further place priors on different categories based on expected location (Supplementary Notes), which is harder to do for real number directional vectors. **b**, Comparison between the SCS model and the SCS model without the transformer attention layers (SCS MLP). Specifically, we removed the attention layer from each of the encoder layers, leaving a two-layer fully connected network (MLP), layer normalization, and residual connections. The concatenation of the projections of neighboring spots ($r$) is fed into the encoder layer. Evaluation on the Stereo-seq dataset shows that SCS achieved an average correlation that was significantly higher than that of SCS MLP (0.60 vs 0.57, Wilcoxon signed-rank test, one-sided; $N$=38,109 cells). **a-b**, Boxplots show medians (horizontal line in each box), interquartile ranges (boxes), 1.5 interquartile (whiskers), and remaining point individuals.

#### Supplementary Tables

**Supplementary Table 1: Runtime and memory usage analysis for different methods on the Stereo-seq dataset.** Due to the environment requirements, Baysor was run on a desktop with Intel Xeon W-2123 @ 3.60GHz CPUs, while other methods were run on a cluster with AMD Opteron 4386 CPUs. The average runtime and memory usage for finished patches of each method are reported.

| Methods | Memory | Time | Number of CPU cores used | Use of GPU |
|---|---|---|---|---|
| SCS | 43.6 GB | 1 hr 4 min 43 s | 8 | Yes |
| Baysor | 8.7 GB | 14 min 1 s | 8 | No |
| JSTA | 18.8 GB | 1 hr 52 min 8 s | 8 | No |
| Watershed | 0.9 GB | 1 min 25 s | 1 | No |
| Cellpose | 1.3 GB | 2 min | 1 | No |
| DeepCell | 1.8 GB | 1 min 56 s | 1 | No |
| StarDist | 1.1 GB | 1 min 33 s | 1 | No |

### References

[1] Ao Chen et al. “Spatiotemporal transcriptomic atlas of mouse organogenesis using DNA nanoball-patterned arrays”. In: *Cell* 185.10 (2022), pp. 1777–1792.

[2] Uwe Schmidt et al. “Cell detection with star-convex polygons”. In: *International Conference on Medical Image Computing and Computer-Assisted Intervention*. Springer. 2018, pp. 265–273.

[3] Carsen Stringer et al. “Cellpose: a generalist algorithm for cellular segmentation”. In: *Nature methods* 18.1 (2021), pp. 100–106.

[4] Dylan Bannon et al. “DeepCell Kiosk: scaling deep learning–enabled cellular image analysis with Kubernetes”. In: *Nature methods* 18.1 (2021), pp. 43–45.

[5] Noah F Greenwald et al. “Whole-cell segmentation of tissue images with human-level performance using large-scale data annotation and deep learning”. In: *Nature biotechnology* 40.4 (2022), pp. 555–565.

[6] Martin Guilliams et al. “Spatial proteogenomics reveals distinct and evolutionarily conserved hepatic macrophage niches”. In: *Cell* 185.2 (2022), pp. 379–396.

[7] F Alexander Wolf, Philipp Angerer, and Fabian J Theis. “SCANPY: large-scale single-cell gene expression data analysis”. In: *Genome biology* 19.1 (2018), pp. 1–5.

[8] Amit Zeisel et al. “Molecular architecture of the mouse nervous system”. In: *Cell* 174.4 (2018), pp. 999–1014.

[9] Chee-Huat Linus Eng et al. “Transcriptome-scale super-resolved imaging in tissues by RNA seqFISH+”. In: *Nature* 568.7751 (2019), pp. 235–239.

[10] Rongxin Fang et al. “Conservation and divergence of cortical cell organization in human and mouse revealed by MERFISH”. In: *Science* 377.6601 (2022), pp. 56–62.

[11] Theodore James Thibault Heiser, Mari-Liis Allikivi, and Meelis Kull. “Shift happens: Adjusting classifiers”. In: *Joint European Conference on Machine Learning and Knowledge Discovery in Databases*. Springer. 2019, pp. 55–70.

[12] Joshua P Gilman, Maria Medalla, and Jennifer I Luebke. “Area-specific features of pyramidal neurons—a comparative study in mouse and rhesus monkey”. In: *Cerebral Cortex* 27.3 (2017), pp. 2078–2094.

[13] Khan L Junatas et al. “Stereological analysis of size and density of hepatocytes in the porcine liver”. In: *Journal of Anatomy* 230.4 (2017), pp. 575–588.
